# Two residues determine nicotinic acetylcholine receptor requirement for RIC-3

**DOI:** 10.1101/2023.01.22.525060

**Authors:** Jennifer D. Noonan, Robin N. Beech

## Abstract

Nicotinic acetylcholine receptors (N-AChRs) mediate fast synaptic signalling and are members of the pentameric ligand-gated ion channel (pLGIC) family. They rely on a network of accessory proteins *in vivo* for correct formation and transport to the cell surface. RIC-3 is an endoplasmic reticulum protein that physically interacts with nascent pLGIC subunits and promotes their oligomerization. It is not known why some N-AChRs require RIC-3 in heterologous expression systems, while others do not. Previously we reported that the ACR-16 N-AChR from the parasitic nematode *Dracunculus medinensis* does not require RIC-3 in *Xenopus laevis* oocytes. This is unusual because all other nematode ACR-16, like the closely related *Ascaris suum* ACR-16, require RIC-3. Their high sequence similarity limits the number of amino acids that may be responsible, and the goal of this study was to identify them. A series of chimeras and point mutations between *A. suum and D. medinensis* ACR-16, followed by functional characterization with electrophysiology, identified two residues that account for a majority of the receptor requirement for RIC-3. ACR-16 with R/K159 in the cys-loop and I504 in the C-terminal tail did not require RIC-3 for functional expression. Mutating either of these to R/K159E or I504T, residues found in other nematode ACR-16, conferred a RIC-3 requirement. Our results agree with previous studies showing that these regions interact and are involved in receptor synthesis. Although it is currently unclear what precise mechanism they regulate, these residues may be critical during specific subunit folding and/or assembly cascades that RIC-3 may promote.

## Introduction

The pentameric ligand-gated ion channels (pLGICs) represent an ancient protein superfamily with functions ranging from neuronal communication in animals ^1^ to acid sensing in bacteria ^2^. Despite an origin predating multicellularism, receptor structure shows remarkable conservation ^3^. Receptors are composed of five structurally similar subunits that form an ion-permeable pore, mediating fast and specific ion flux across membranes ^4, 5^. A large extracellular domain (ECD) binds the activating ligand and the four transmembrane domains (TM1 - 4) of each subunit anchor the receptor in the membrane and form the central ion pore ^6, 7^. Eukaryotic pLGICs are known as cys-loop receptors because they contain a cys-loop motif of 13 amino acids between two cysteines that form a disulphide bridge in the ECD that is critical for receptor gating ^3, 6^. A highly variable intracellular loop (ICL) in the eukaryotic receptors remains poorly characterized for structure and function ^3, 8^. Finally, a short C-terminal tail protrudes into the extracellular space directly after the last transmembrane domain (TM4) that has been implicated in receptor gating ^9^.

During receptor synthesis, subunits undergo large and specific structural changes ensuring appropriate folding and oligomerization as monomers combine into a mature pentamer ^10, 11^. A majority of the expression signals are found within the receptor ECD and TM1 ^11–15^, however the ICL and C-terminal tail contribute as well ^16–23^. This process is tightly regulated by a variety of accessory proteins that function as chaperones, mediating subunit assembly ^24^, thiol redox reactions ^25^, glycosylation ^26, 27^ and trafficking^28^ . Defects in these accessory proteins may be associated with genetic disease and altered behaviour ^10, 29^. The receptor expression machinery itself is complex, with hundreds of proteins involved and where only a fraction have been characterized ^24, 30^. Additionally, even closely related receptor subunits can be differentially regulated by the same accessory protein ^31, 32^. Although chaperones continue to be identified ^25^, it remains unclear why some receptors require specific accessory proteins to facilitate functional synthesis while others do not.

Resistance to cholinesterase 3 (RIC-3) is one of the most well-known accessory proteins involved in production of the acetylcholine receptor (AChR) family ^33^. RIC-3 also regulates serotonin receptor (5-HT_3_R) expression ^34^ however anionic glycine, glutamate and γ-aminobutyric acid (GABA) receptor expression is not affected by it ^35, 36^. RIC-3 was first identified in the model nematode *Caenorhabditis elegans* and has since been found to have versatile effects on both vertebrate and invertebrate receptors alike ^37, 38^. In general, it promotes the expression of AChRs while inhibiting 5-HT_3_Rs ^34, 39–41^. However, there are examples that deviate from this general trend ^32, 42, 43^. In fact, expression of specific receptors from combinations with similar subunit compositions can be arrested during transport to enrich a specific receptor subtype at the cell surface ^44^. RIC-3 functions in the endoplasmic reticulum (ER) by promoting subunit assembly, and in its absence subunits remain in the ER unassembled ^45, 46^. Sequence analysis predicts it has 1 or 2 N-terminal transmembrane domains with an intracellular coiled-coil C-terminal domain ^47^. Both of these regions physically interact with and are required for stabilizing immature nematode AChR subunits ^32, 43^. Although functional regions within RIC-3 have been identified ^32, 34, 43, 45, 48^, few have studies have investigated regions of the receptors that may interact with or confer a RIC-3 requirement. The role of RIC-3 in production of the mouse 5-HT_3_R has been shown to depend on the presence of the 5-HT_3_R ICL, and more specifically, interacts with the first 24 amino acids of the IC L ^49, 50^. RIC-3 interacts with subunits, but it is not known what exact mechanism it regulates when promoting subunit folding/oligomerization.

The acetylcholine receptor subunit 16 (ACR-16) subunit of nematodes produces a homomeric AChR that responds specifically to nicotine and regulates the contraction of body muscle ^51, 52^. The ACR-16 nicotinic acetylcholine receptors (N-AChRs) of several different nematode species, including the human parasite, *Ascaris suum* (Asu-ACR-16), have been characterized *ex vivo* in the *Xenopus laevis* oocyte expression system where they require the co-injection of RIC-3 ^53–59^. This expression system allows for transient and specific expression of injected complimentary RNA (cRNA) and characterization of receptors in a controlled environment ^60^. Previously, we reported that a robust N-AChR from the human parasitic nematode *Dracunculus medinensis* (Dme-ACR-16), that is closely related to *A. suum*, can be expressed in *X. laevis* oocytes in the absence of RIC-3 ^61^. Addition of RIC-3 led to a higher Dme-ACR-16 response that was similar to that from Asu-ACR-16, suggesting that there is still an interaction that promotes receptor production but that the requirement is no longer indispensable for Dme-ACR-16.

The sequence identity between Asu-ACR-16 and Dme-ACR-16 is 84%, we thus hypothesized that relatively few positions determine the RIC-3 requirement and that they can be identified using these phylogenetically related receptors with differing RIC-3 dependencies. We therefore sought to identify the sequences responsible using a series of chimeras and point mutations between *A. suum* and *D. medinensis* ACR-16, followed by functional measurements using electrophysiology. Knowing molecular determinants of accessory protein requirement and may shed light on the specific functions that RIC-3 regulates during subunit folding and assembly. We identified two residues that have removed the requirement for RIC-3 in Dme-ACR-16. When the receptors had both the cys-loop K/R159 and the C-terminal tail I504, they no longer required RIC-3 for function. Mutating either residue to amino acids found in other nematode ACR-16 (K/R159E or I504T) abolished responses without RIC-3. Their proximity to one another in the subunit structure suggests that they may participate in critical interactions during subunit production, obviating the need for RIC-3. Molecular dynamic simulations confirmed the residues are close enough to interact but were unable to explain the difference in requirement for RIC-3. This may be because our simulations were performed on models based on mature subunit structure templates that have already gone through the structural changes necessary for producing a functional receptor. Nevertheless, our results agree with previous work that found that the cys-loop and C-terminal tail directly interact ^9^ and that they are involved in receptor expression ^62^. Here we identified two residues that are critical in regulating AChR functional synthesis and provides novel insight into a possible molecular mechanism that determines RIC-3 requirement.

## Results

### *D. medinensis* ACR-16 does not require RIC-3 for heterologous function

We previously reported that *D. medinensis* ACR-16 no longer required RIC-3 co-injection for function in the oocyte expression system, whereas the closely related *A. suum* ACR-16 did ^61^. Their high sequence identity (84%) provided an opportunity to identify the sequence responsible (Figure S1A). The coding sequences of *A. suum* and *D. medinensis* ACR-16 were modified to introduce complimentary restriction enzyme recognition sites to facilitate sequence exchange in a way that did not alter the amino acid sequence (referred to as ACR-16-REMs, Figure 1D-G, Figure S1A). Nomenclature for the chimeras indicates the region that has been exchanged, for example Asu-dmeECD indicates the ECD has been removed from the *A. suum* receptor and replaced with the *D. medinensis* ECD.

**Figure 1.**
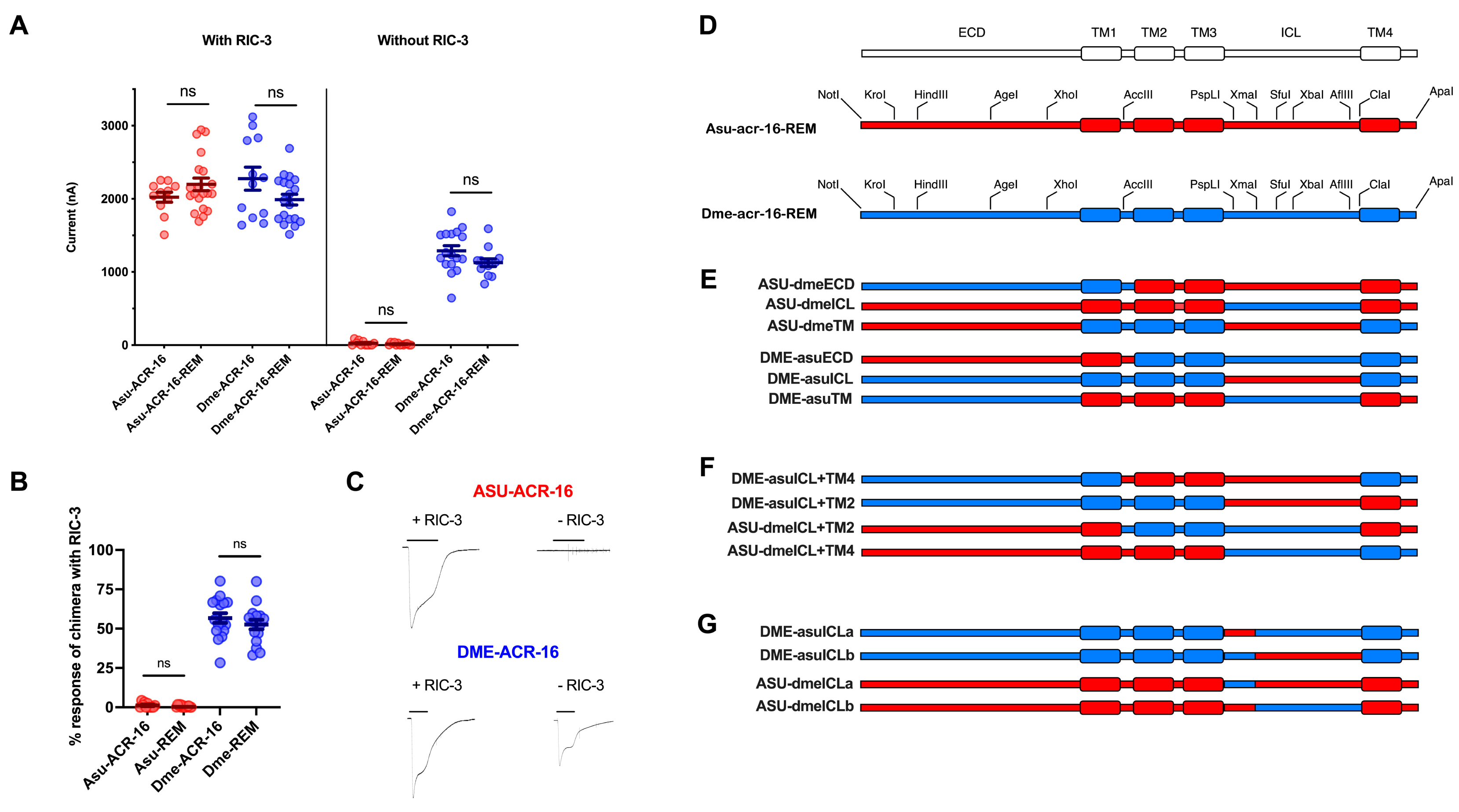
*D. medinensis* ACR-16 does not require the RIC-3 accessory protein for function in oocytes. *A. suum* ACR-16 (Asu-ACR-16) and *D. medinensis* ACR-16 (Dme-ACR-16) receptors have different requirements for the RIC-3 accessory protein. Their high sequence similarity indicates that few regions determine this and that they can be identified using receptor chimeras. **(A)** Restriction enzyme modified sequences (labelled ACR-16-REM) produce responses indistinguishable from their original sequences. *A. suum* ACR-16-REM and *D. medinensis* ACR-16-REM produce similar response with RIC-3 (2197 ± 86 nA, p = 0.1798 for Asu-ACR-16 and 1989 ± 73 nA, p = 0.0732 for Dme-ACR-16) whereas current without RIC-3 is minimal for Asu-ACR-16-REM (14 ± 4 nA, p = 0.3265) and large for Dme-ACR-16-REM (1127 ± 53 nA, p = 0.0891). Any change in chimera response without RIC-3 would be due to the exchanged region. Error bars represent standard error. Responses of the restriction enzyme modified constructs are compared to those of native receptor sequence previously reported ^61^. **(B)** Current responses without RIC-3 shown as a percent of response with RIC-3 for Asu-ACR-16 (1.19 ± 0.49 %) compared to Asu-ACR-16-REM (0.62 ± 0.2 %, p = 0.2627) and Dme-ACR-16 (56.7 ± 3 %) compared to Dme-ACR-16-REM (52.6 ± 3 %, p = 0.3552). Error bars represent standard error. **(C)** Example recordings of current responses are shown with and without RIC-3 for Asu-ACR-16 (top) and Dme-ACR-16 (bottom). **(D)** Chimeras were made to identify the regions contributing to RIC-3 requirement in ACR-16. *A. suum* and *D. medinensis* coding sequences were modified to introduce complementary restriction enzyme sites (ACR-16-REM) to generate the chimeras shown in **(E-G).** Asu-ACR-16 sequence in red, Dme-ACR-16 sequence in blue. **(E)** Chimeras exchanging the three main structural regions ECD, TM and ICL. **(F)** Chimeras exchanging each TM2 or TM4 along with the ICL. **(G)** Chimeras cutting the ICL in half with either *D. medinensis* or *A. suum* TM4.

Receptor current in response to acetylcholine (ACh) and RIC-3 requirement from these engineered sequences (ACR-16-REM) were indistinguishable from their unmodified counterparts (Figure 1A) ^61^. Therefore, any changes observed in responses without RIC-3 in the chimeras would be due to the exchanged region of interest. Compared to previous reports ^61^, responses for Asu-ACR-16-REM with RIC-3 were 2197 ± 84 nA (p=0.1798) and without RIC-3 were 14 ± 4 nA (p=0.3265). While responses for Dme-ACR-16-REM with RIC-3 were 1989 ± 73 nA (p=0.0732) and without RIC-3 were 1127 ± 53 nA (p=0.0891). Dme-ACR-16 did not require RIC-3 to produce robust responses in oocytes. In the absence of RIC-3, Dme-ACR-16-REM produces current sizes of 52.6 ± 3% compared to sizes when RIC-3 was included. In contrast, Asu-ACR-16-REM response without RIC-3 was only 0.6% compared to sizes when RIC-3 is included (Figure 1B). We next wanted to identify the main structural regions mediating this difference.

### The intracellular loop, transmembrane domain regions and C-terminal tail determine RIC-3 requirement

Chimeras were first made by exchanging each one of the three main structural domains (ECD, TMs and ICL, Figure 1E) followed by smaller exchanges to narrow down the sequence of interest (Figure 1G & H). RIC-3 co-injection with each chimera served as a control to ensure the chimera in question was functional. All chimeras were functional however some showed significantly higher or lower responses compared to their reference receptor even when including RIC-3 (Figure S2A). Therefore we show chimera response as a percent without RIC-3 compared to with RIC-3 to control for any differences in response due to the chimera construct. The ICL, TMs and C-terminal tail were found to contribute to RIC-3 requirement. Percent responses for the reference receptors Asu-ACR-16-REM and Dme-ACR-16-REM were 0.6 ± 0.2% and 52.6 ± 3%, respectively (Figure 1B). The requirement for RIC-3 was not affected by exchanging the ECD (Figure 2A). The Asu-ACR-16 with the *D. medinensis* ECD chimera (Asu-dmeECD) was unable to produce responses without RIC-3 (0 ± 0%, p<0.01), while the Dme-ACR-16 with the *A. suum* ECD chimera (Dme-asuECD) produced large currents in the absence of RIC-3 (69 ± 5%, p<0.01). Exchange of the ICL reversed some, but not all, of the RIC-3 requirement. Both chimeras responded minimally without RIC-3 (14.9 ± 2%, p<0.0001 for Asu-dmeICL and 11.3 ± 5%, p<0.0001 for Dme-asuICL). The TMs and C-terminal exchanges nearly completely reversed the RIC-3 requirement. The Asu-ACR-16 with the *D. medinensis* TMs chimera (and C-terminal tail) had large responses (Asu-dmeTM, 39 ± 3%, p<0.0001) compared to those of the Asu-ACR-16-REM. The Dme-ACR-16 with the *A. suum* transmembrane domains chimera (and C-terminal tail) now had reduced responses (Dme-asuTM, 2.0 ± 0.4%, p<0.0001) compared to Dme-ACR-16-REM. These chimeras indicate that the ICL, TMs and C-terminal tail determined AChR requirement for the RIC-3 accessory protein, with the TMs and C-terminal tail having the most effect.

**Figure 2.**
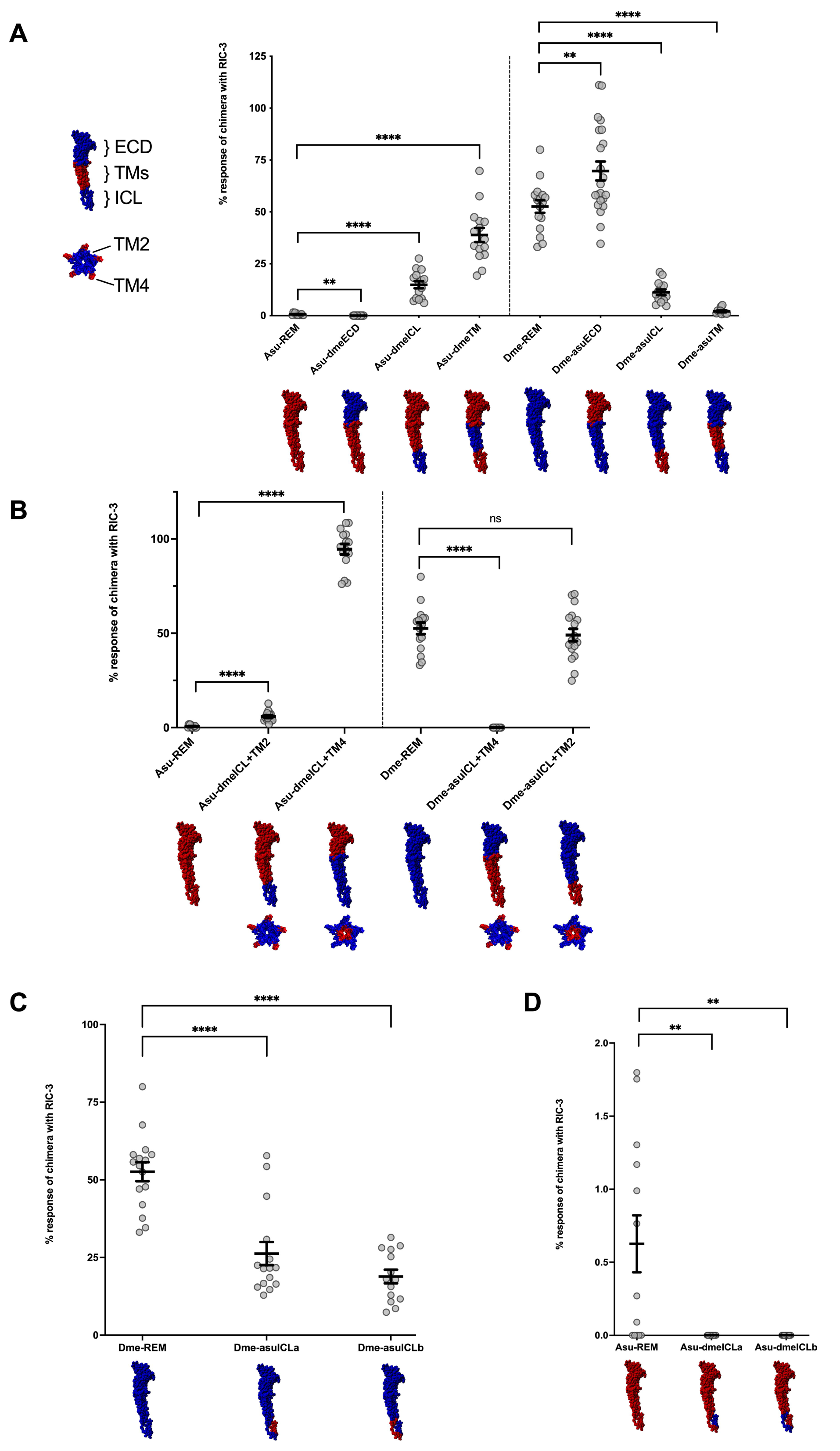
Main structural region chimeras show the intracellular loop, transmembrane domain 4 and C-terminal tail determine RIC-3 requirement. **(A)** Chimeras were made exchanging each of the three main structural regions (ECD, ICL or TM) to identify which regions mediate RIC-3 requirement. Percent response of the chimeras with RIC-3 compared to response without RIC-3 are shown. Asu-ACR-16-REM responses increased upon addition of the *D. medinensis* ICL (Asu-dmeICL, 14.98 ± 1.7%, p<0.0001) or the TM + C-terminal tail (Asu-dmeTM, 38.88 ± 3.4, p<0.0001) and decreased with the *D. medinensis* ECD (Asu-dmeECD, 0 ± 0%, p<0.01). Dme-ACR-16-REM responses decreased upon addition of the *A. suum* ICL (Dme-asuICL, 11.25 ± 1.4%, p<0.0001) or the TM + C-terminal tail (Dme-asuTM, 2.05 ± 0.4%, p<0.0001) and increased with addition of *A. suum* ECD (Dme-asuECD; 69.7 ± 4.7%, p<0.01). **(B)** Substitutions in the transmembrane domains are only found in TM2 and TM4, therefore chimeras were made exchanging individually TM2 or TM4 along with the ICL to identify if one or both of those transmembrane domains mediate RIC-3 requirement. Percent response of the chimeras without RIC-3 compared to responses with RIC-3 are shown. Asu-ACR-16-REM with the *D. medinensis* ICL and TM2 (Asu-dmeICL+TM2, 5.9 ± 0.7%, p<0.0001) and the *D. medinensis* ICL and TM4 (Asu-dmeICL+TM4, 94.56 ± 2.8%, p<0.0001) produced larger percent responses, with the Asu-dmeICL+TM4 chimera having the highest response compared to Asu-dmeICL+TM2 (p<0.0001). Asu-ACR-16 with the *A. suum* ICL and TM2 did not affect percent response (Dme-asuICL+TM2, 49.1 ± 3.3%, p=0.4400) whereas addition of the *A. suum* ICL and TM4 inhibited all percent responses (Dme-asuICL+TM4, 0.0 ± 0%, p<0.0001). Therefore the TM4 and C-terminal, and not the TM2, determine RIC-3 requirement. **(C)** Intracellular loop chimeras in the *D. medinensis* receptor have responses lower than the *D. medinensis* receptor indicating both halves contribute to RIC-3 requirement. Dme-asuICLa (26.27 ± 3.7%, p<0.0001) and Dme-asuICLb (18.88 ± 2.2%, p<0.0001). **(D)** Intracellular loop chimeras in the *A. suum* receptor have no responses without RIC-3 indicating both halves are needed together for RIC-3 requirement. Asu-dmeICLa (0 ± 0%, p<0.01) and Asu-dmeICLb (0 ± 0 %, p<0.01). Subunit structures show chimera sequences with red regions indicating *A. suum* sequence and blue regions indicating *D. medinensis* sequence. Error bars represent standard error. **p<0.01; ****p<0.0001.

### The fourth transmembrane domain and the C-terminal tail determine RIC-3 requirement

*A. suum* and *D. medinensis* ACR-16 TM sequences are highly conserved, yet these regions accounted for the majority of the RIC-3 requirement. Substitutions are only observed in the second (TM2) and fourth (TM4) transmembrane helices and only one in the C-terminal tail (Figure S1B). Chimeras exchanging TM2 or TM4 were made to identify if one or both TMs contribute to the requirement for RIC-3 (Figure 1F). Two pairs of these chimeras were made; with and without the *D. medinensis* ACR-16 ICL to measure the effect of each transmembrane domain individually, and in conjunction with the ICL since It was found to also contribute. The TM2 did not contribute to RIC-3 requirement based on the chimeras that had the *D. medinensis* ACR-16 TM2 (5.8 ± 0.7%, p<0.0001 for Asu-dmeICL+TM2 & 0 ± 0% p<0.0001 for Dme-asuICL+TM4, Figure 2B). In contrast, the TM4 and the C-terminal mediated RIC-3 requirement based on the chimeras that had these *D. medinensis* ACR-16 regions (94 ± 3%, p<0.0001 for Asu-dmeICL+TM4 & 49 ± 3%, p=0.4400 for Dme-asuICL+TM2, Figure 2B). These chimeras ruled out the possibility that TM2 is involved and indicates that it is entirely within the TM4 the C-terminal tail sequences.

### The entire ICL contributes to RIC-3 requirement

The ICL contributed minimally to receptor requirement for RIC-3 (Figure 2A). Narrowing down the specific ICL sequence(s) that regulate RIC-3 requirement would not be straightforward because the ICL sequence is highly variable and most of its structure remains undetermined ^8^. However, the first 24 amino acids of a mouse 5-HT_3_R ICL have been found to interact with RIC-3 ^50^. Due to the longer length of AChR ICLs, sequence alignment indicates that these 24 amino acids in a 5-HT_3_R correspond to the first 50 amino acids in these AChRs and provided us with a reference with which to compare to. The design of our REM sequences allowed us to exchange first 62 amino acids of the ICL, constructed under both *A. suum* and *D. medinensis* ACR-16 TM4 + C-terminal tail (Figure 1G).

Neither half of the ICL contained the region that confers RIC-3 requirement. Chimeras that split the *D. medinensis* ACR-16 ICL were compared to Dme-ACR-16 (Figure 2C). If the region was confined to only one of these halves, then it would be expected to have response equal to Dme-ACR-16. Both chimeras produced responses that were significantly smaller than Dme-ACR-16 (Dme-asuICLa 26 ± 4%, p<0.0001 & Dme-asuICLb 18 ± 2%, p<0.0001). This was further confirmed with the second pair of chimeras that split the ICL in half in the *A. suum* ACR-16 receptor (Figure 2D). The requirement for RIC-3 could not be conferred by either segment of the ICL alone. Percent responses were smaller compared to Asu-ACR-16 (Asu-dmeICLa 0 ± 0%, p<0.01, Asu-dmeICLb 0 ± 0%, p<0.01), suggesting the requirement was distributed across both regions. Therefore, the sequence within the ICL that mediates an AChR requirement for RIC-3 is not confined to the first half, unlike 5-HT_3_Rs ^50^.

### The C-terminal tail determines RIC-3 requirement

The TM4 and C-terminal tail were found to determine a majority of the RIC-3 requirement (Figure 2B). There are three amino acid substitutions between the *A. suum* and *D. medinensis* ACR-16 TM4 and one in the C-terminal tail. These are residues 485, 488, and 491 in TM4 and 504 in the C-terminal tail (as numbered in the alignment in Figure S1B). Dme-ACR-16 has isoleucine at all these positions, whereas Asu-ACR-16 has three valines in the TM4 and a threonine in the C-terminal tail. To determine the combination of these substitutions that contribute to the effect, a series of point mutations were made mutating the sites in *A. suum* to those in *D. medinensis*, and vice versa (Figure 3A & C). Point mutation nomenclature TM4-1 refers to residue 485, TM4-2 refers to residue 488, TM4-3 refers to residue 491, and tail refers to residue 504. All TM4 and C-terminal tail point mutations produced robust responses with RIC-3 with some significant deviations from their reference receptor responses (Figure S3A & S3B).

**Figure 3.**
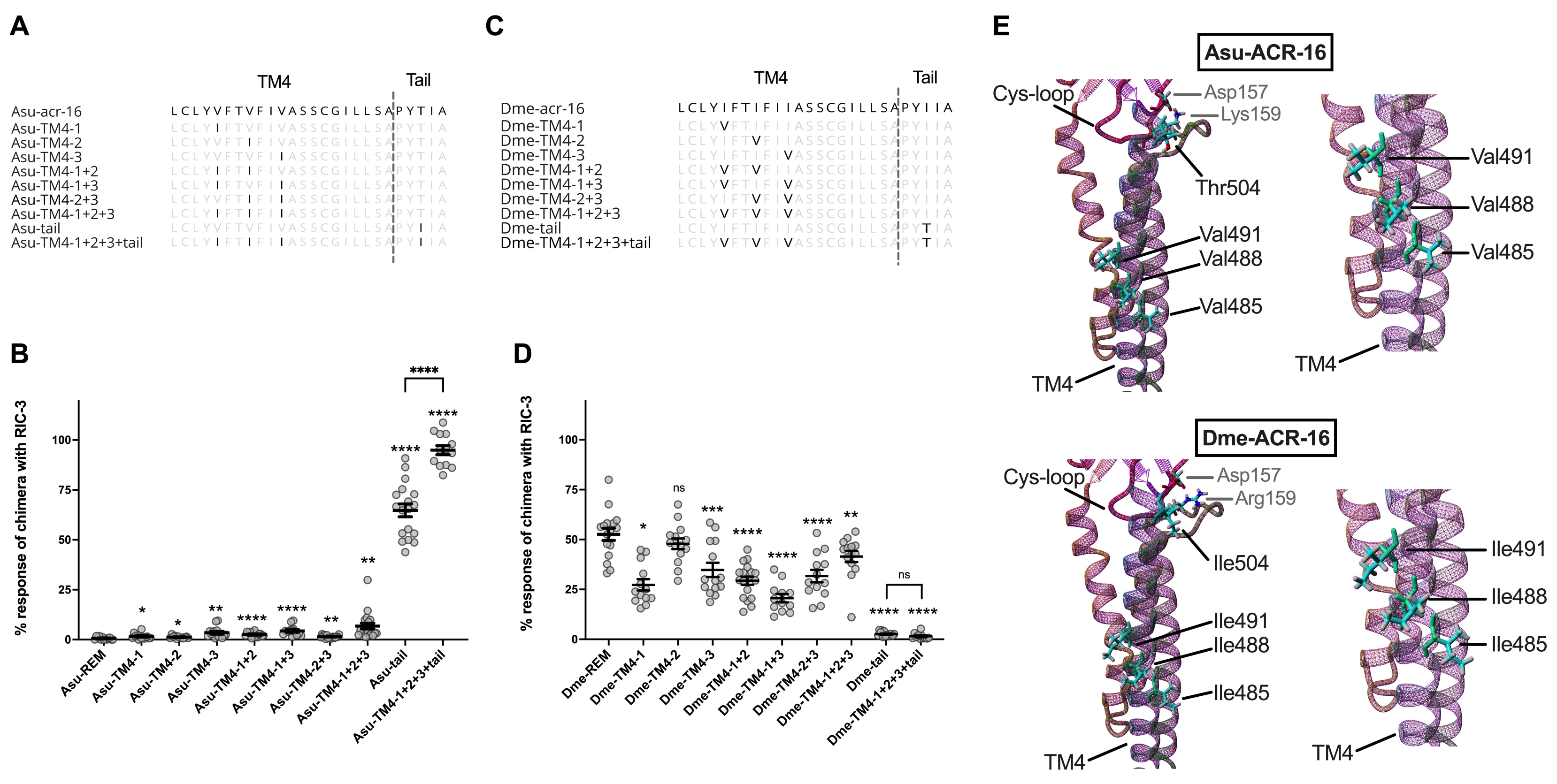
A single amino acid in the C-terminal tail determines RIC-3 requirement in ACR-16. Point mutations were made in Asu-ACR-16 and Dme-ACR-16 TM4 and the C-terminal tail. Amino acid position 485 refers to TM4-1, 488 refers to TM4-2, 491 refers to TM4-3, 504 refers to tail. **(A)** Point mutations in Asu-ACR-16 shown in bold. **(B)** Point mutations in Dme-ACR-16 shown in bold. **(C)** Percent response of Asu-ACR-16 point mutations of response without RIC-3 compared to response with RIC-3. The single, double and triple TM4 point mutations marginally increased responses; Asu-TM4-1 (1.6 ± 0.4%, p<0.05), Asu-TM4-2 (1.2 ± 0.2%, p<0.05), Asu-TM4-3 (3.4 ± 0.7, p<0.01), Asu-TM4-1+2 (2.6 ± 0.3%, p<0.0001), Asu-TM4-1+3 (4.3 ± 0.7%, p<0.0001), Asu-TM4-2+3 (1.4 ± 0.2%, p<0.01) and Asu-TM4-1+2+3 (6.7 ± 1.5%, p<0.01). The single point mutation to the tail alone was enough to produce large response without (Asu-tail, 64.73 ± 3.2 %, p<0.0001), and this was increased with the addition of the three TM4 mutations (Asu-TM4-1+2+3+tail (94.9 ± 2.3 %, p<0.0001). **(D)** Percent response of Dme-ACR-16 point mutations of response without RIC-3 compared to response with RIC-3. Except for Dme-TM4-2 (47.8 ± 2.7%, p=0.2551), all single double and triple point mutations lowered responses without RIC-3, but did not abolish them; Dme-TM4-1 (27.3 ± 3.8%, p<0.0001), Dme-TM4-3 (34.8 ± 3.6%, p<0.005), Dme-TM4-1+2 (29.4 ± 2.0%, p<0.0001), Dme-TM4-1+3 (20.6 ± 2.1%, p<0.0001), Dme-TM4-2+3 (31.7 ± 3.1%, p<0.0001), Dme-TM4-1+2+3 (41.5 ± 2.8%, p<0.05). The single point mutation to the tail abolished response without RIC-3 (Dme-tail, 2.6 ± 0.3%, p<0.0001), and addition of the three TM4 mutations (Dme-TM4-1+2+3+tail,1.5 ± 0.5%, p<0.0001) produced similar response. These point mutation constructs indicate the single C-terminal tail residue determines RIC-3 requirement. Mutating T504I in Asu-ACR-16 no longer needed RIC-3, while I504T mutation in Dme-ACR-16 needed RIC-3, therefore the tail residue determines RIC-3 requirement in the oocyte expression system. **(E)** All four residues are aligned on the outside of the subunit. Error bars represent standard error. *p<0.05; **p<0.01; ***p<0.0005; ****p<0.0001.

When comparing percent response, all single, double, and triple point mutations in the *A. suum* TM4 (VàI) had small responses without RIC-3; Asu-TM4-1 (1.6 ± 0.4%, p<0.05), Asu-TM4-2 (1.2 ± 0.2%, p<0.05), Asu-TM4-3 (3.4 ± 0.7%, p<0.01), Asu-TM4-1+2 (2.6 ± 0.3%, p<0.0001), Asu-TM4-1+3 (4.3 ± 0.7%, p<0.0001), Asu-TM4-2+3 (1.4 ± 0.2%, p<0.01) and Asu-TM4-1+2+3 (6.7 ± 1.5%, p<0.01) (Figure 3B). The C-terminal tail mutation on its own (T504I) produced large response without RIC-3 (Asu-tail; 64 ± 4%, p<0.0001). Interestingly, the C-terminal tail mutation in combination with the three TM4 mutations produced even greater responses without RIC-3 (Asu-TM4-1+2+3+tail; 95 ± 3%, p<0.0001). Therefore, I504 determines RIC-3 requirement and the three TM4 sites contribute as well.

These *A. suum* point mutations would predict that all of the single, double and triple point mutations of the *D. medinensis* TM4 region would function with responses slightly below those of the native sequence without RIC-3, and that the two tail mutant conditions would induce the lowest responses. This was indeed observed (Figure 3D). The C-terminal tail mutation on its own (I504T) nearly abolished all response without RIC-3 (Dme-tail; 2.6 ± 0.3%, p<0.0001), as did the tail in combination with the three TM4 mutations (Dme-TM4-1+2+3+tail; 1.5 ± 0.5%, p<0.0001). Except for Dme-ACR-16-TM2 (47.8 ± 2.7%, p=0.2551) which did not produce any changes to relative percent response without RIC-3, the TM4 point mutations (IàV) produced responses that were significantly smaller than Dme-ACR-16 but not as small as the tail mutation conditions; Dme-TM4-1 (27.3 ± 3.8%, p<0.0001), Dme-TM4-3 (34.8 ± 3.6%, p<0.005), Dme-TM4-1+2 (29.4 ± 2.0%, p<0.0001), Dme-TM4-1+3 (20.6 ± 2.1%, p<0.0001), Dme-TM4-2+3 (31.7 ± 3.1%, p<0.0001), Dme-TM4-1+2+3 (41.5 ±2.8%, p<0.05).

### A basic cys-loop residue mediates RIC-3 requirement

Our results indicate that I504 alleviates the requirement for RIC-3. Therefore, we would predict that other AChRs that also contain this isoleucine would not need RIC-3. Curiously, ACR-16 from related nematodes, that also have I504 still require RIC-3 co-injection ^61^ which led us to believe that another region may interact with the tail, and together they determine RIC-3 requirement. The C-terminal tail directly follows the TM4 and protrudes into the extracellular space. It has been found to interact with residues within the cys-loop motif in the ECD ^9, 63^, and the cys-loop has been found to undergo structural changes essential for subunit oligomerization ^11^. We therefore considered the idea that the C-terminal tail and cys-loop may participate in interactions that could promote subunit folding/assembly thus no longer requiring RIC-3.

An alignment of nematode ACR-16 that have been previously characterized in the oocyte system identified two positions in the cys-loop motif with substitutions, positions 157 and 159 in Figure 4A. *Gonglyonema pulchrum* is a parasitic nematode phylogenetically related to both *A. suum* and *D. medinensis,* and although its ACR-16 sequence contains I504, it requires RIC-3 for function^61^. The *G. pulchrum* cys-loop substitutions were therefore used to investigate a potential interaction between the cys-loop and tail for RIC-3 requirement. The two cys-loop substitutions, together with the C-terminal tail position 504, in *A. suum* ACR-16 were D157-K159-T504, in *D. medinensis* ACR-16 were D157-R159-I504 and in *G. pulchrum* ACR-16 were N157-E159-I504 (Figure 4A). We next mutated the two cys-loop residues individually or together in the *D. medinensis* ACR-16 sequence to those found in *G. pulchrum* ACR-16. All three mutation combinations produced lower responses without RIC-3 (Figure 4C & D). The Dme-D157N substitution did reduce percent responses (17.39 ± 1.9%, p<0.0001) but not as much as the Dme-R159E substitution (0.55 ± 0.16%, p<0.0001). Combining both cys-loop substitutions together, Dme-D157N+R159E, had minimal percent responses (0.06 ± 0.03%, p<0.0001). These results identified both residues contribute to the RIC-3 requirement, however the R159 can completely determine it.

**Figure 4.**
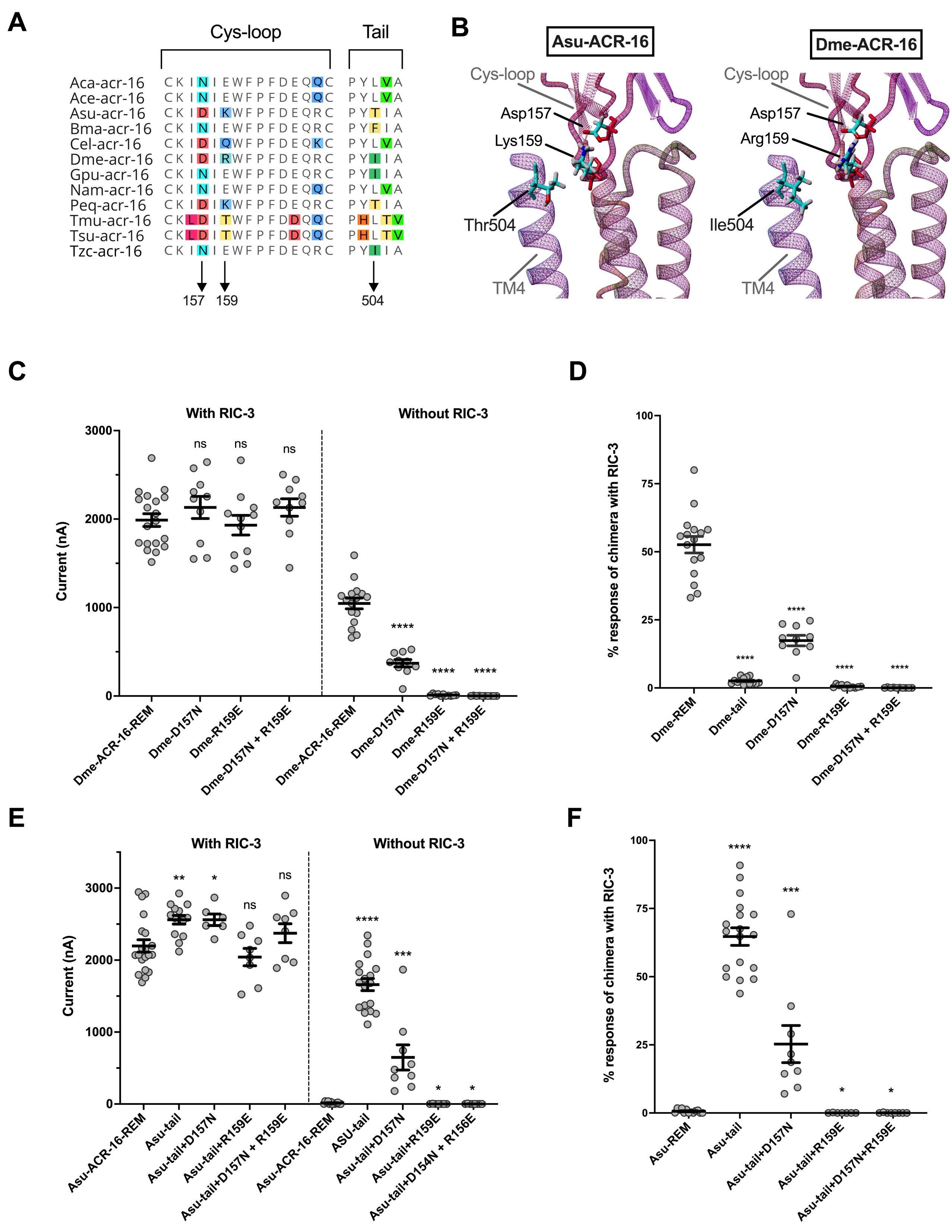
Residues in the cys-loop and C-terminal tail determine RIC-3 requirement. The observation that other ACR-16 receptors not studied here require RIC-3 even though they have the I504 residue indicates some other region, together with the tail, contributes to the effect. The ECD cys-loop is known to interact with the tail ^9, 63^, and both regions are involved in receptor expression ^11^ therefore we hypothesized that residues within the cys-loop may interact with I504 to mediate the RIC-3 requirement. **(A)** alignment of all characterized nematode ACR-16 cys-loop and tail sequences identified two substitutions of interest in the cys-loop (157 and 159) and within proximity to I504 in the subunit structure **(B)**. **(C)** Single and double cys-loop point mutations were made in the *D. medinensis* sequence to the residues found in the other ACR-16. All three cys-loop mutation conditions produced responses indistinguishable from those of the *D. medinensis* receptor when RIC-3 was included, Dme-D157N (2131 ± 125 nA, p=0.2983), Dme-R159E (1930 ± 111 nA, p=0.6507) and Dme-D157N + R159E (2131 ± 99 nA, p=0.2571). However, significantly lower responses were measured when RIC-3 was omitted under all three conditions; Dme-D157N produced lower, but measurable responses (370.5 ± 41.2nA, p<0.0001), Dme-R159E produced smaller currents (10.55 ± 13.3 nA, p<0.0001), and the double cys-loop mutation Dme-D157N + R159E produced minimal to no response (1.2 ± 0.7 nA, p<0.0001). **(D)** When shown as a percent response, all cys-loop mutations produced severely inhibited responses; Dme-D157N (17.39 ± 1.9%, p < 0.0001), Dme-R159E (0.54 ± 0.16%, p < 0.0001) and Dme-D157N + R159E (0.06 ± 0.03, p < 0.0001). **(E)** Single and double cys-loop point mutations were made in the *A. suum* + T504I mutation construct (*A. suum* with the T504I mutation, Figure 3A; Asu-tail) to the residues found in the other nematode ACR-16. All three mutation constructs produced functional receptors when RIC-3 was included. Asu-tail+D157N produced higher responses (2560 ± 80.38 nA, p<0.05) while Asu-tail+R159E (2042 ± 120.4 nA, p=0.3294) and the double cys-loop mutations Asu-tail+D157N + R159E (2374 ± 131 nA, p=0.2773) produced currents indistinguishable from the Asu-ACR-16-REM reference. When RIC-3 was omitted; Asu-tail+D157N produced current (647 ± 175 nA, p<0.0005) while both Asu-tail+R159E (0.57+ 0.57 ± 3.13 nA, p<0.05), and the double point mutation Asu-tail+D157N + R159E (0.88 ± 0.88 nA, p<0.05) produced lower responses compared to Asu-ACR-16-REM. **(F)** When shown as a percent response, Asu-tail+D157N (25.29 ± 6.8% p<0.0005) had higher responses and Asu-tail+R159E (0.028 ± 0.028%, p<0.05) and Asu-tail+D157N + R159E (0.037 ± 0.0337%, p<0.05) had lower responses compared to Asu-ACR-16-REM. Error bars represent standard error. *p<0.05; **p<0.01; ***p<0.0005; ****p<0.0001.

To further validate our findings that these cys-loop residues together with I504 determine RIC-3 requirement, we wanted to test if we could introduce a RIC-3 requirement after having removed it. To do so we mutated the same cys-loop residues in the Asu-TM4-tail construct from Figure 3A. This construct has the T504I mutation in the *A. suum* sequence which caused it to no longer require RIC-3 (Figure 3B). All three cys-loop mutation combinations produced lower responses without RIC-3. The Asu-tail+D157N substitution did reduce percent responses (25.25 ± 6.8%, p<0.0001) but not as much as the Asu-tail+R159E substitution (0.03 ± 0.03%, p<0.0001).

Combining both cys-loop substitutions together, Asu-tail+D157N + R159E, again produced barely any responses (0.04 ± 0.04%, p<0.0001). To summarize, we were able to remove the Asu-ACR-16 requirement for RIC-3 by mutating T504I, and this phenotype was subsequently reverse by addition of a second mutation K/R159E. These results validate that the residues in the cys-loop motif and C-terminal tail can determine AChR requirement for RIC-3.

Our point mutations suggest a possible interaction between the C-terminal tail 504 and the cys-loop residues 157, 159. We therefore wanted to see if any interactions between the C-terminal tail and cys-loop would account for the differences in RIC-3 requirement using *in silico* simulations. Molecular dynamic simulations can provide a way to assess molecular interactions if the predicted molecular structures are sufficiently close to reality. Most available template structures for homomeric AChRs lack the tail beyond TM4 ^64, 65^ making a reasonable model of this region problematic. A recent structure for the alpha-7 AChR (PDB: 7KOO) positions the tail as an alpha-helix parallel to the membrane, away from the cys-loop, precluding interaction with the residues in question ^66^. In order to evaluate any interaction between residues 157, 159 and 504 we carried out short simulations (∼10 ns) for a single, membrane-embedded subunit model under the different combinations of substitutions tested here for residues 157, 159 and 504 ^67^. We arbitrarily extended the tail to orient the 504 residue within 3 Å from residues 157 and 159 and minimized the energy of this structure configuration prior to beginning the simulations. Three different simulations, each starting from the same membrane embedded structures, resulted in simulations that were generally similar for each original ACR-16, as well as for all the sequence combinations Asu-ACR-16 D157-E159-I504, D157-K159-I504, N157-E159-I504 and 157N-K159-I504 and Dme-ACR-16 D157-E159-I504, D157-R159-T504, N157-E159-I504 and N157-R159-I504. In no case did interactions between the cys-loop and residue 504 persist for more than a few 25 fs frames of the simulations. The lack of any difference in tail and cys-loop interaction and/or positioning may be because these structures are all mature receptors, whereas RIC-3 is expected to interact with single subunits in the ER membrane when the tertiary structure undergoes significant rearrangement that includes an interaction between the tail and the cys-loop. Regardless, our functional measurements using electrophysiology have identified that residues 159 and 504 can determine AChR requirement for the RIC-3 accessory protein.

## Discussion

Here we identified two residues that determine N-AChR requirement for the RIC-3 accessory protein and by extension, residues that are critical in receptor oocyte expression. The oocyte expression system is an important tool for receptor identification and characterization. The injection of specific subunits and accessory proteins allows us to investigate functional significance in an isolated system. However, if essential components to the production of the channel are not included endogenously by the oocyte or exogenously in the injection mixture, then receptors fail to be expressed. The complex and largely uncharacterized assembly and trafficking of the pLGICs means that it is difficult to determine which component is missing. Many invertebrate AChRs require the co-injection of specific accessory proteins enabling their functional expression ^53, 68^. This is also true for the vertebrate alpha7 receptor ^24^ and appears to be a characteristic of AChRs in general. RIC-3 is one such accessory protein and functions by physically interacting with monomeric subunits, promoting their pentameric oligomerization ^69^.

*A. X. laevis* RIC-3 is expressed in oocytes ^70^, however it might be expressed in too low amounts or be too divergent to promote expression of some receptors ^14^. While many studies have identified the regions in RIC-3 that are required for its function ^32, 43, 71^, little is known about the regions in the receptors themselves that may interact with RIC-3 or why some receptors require it but not others. Previously we reported that the expression of ACR-16 from the human parasite *D. medinensis* is possible in the absence of RIC-3 ^61^. This is in contrast to the closely related pig parasite *A. suum* ACR-16 which requires its co-injection for function ^56, 61^. *A. suum* and *D. medinensis* ACR-16 receptors have similar pharmacology and identical maximal response, yet they have dramatically different requirements for the accessory protein ^61^. Since the co-injected RIC-3 was constant, the cause must be encoded in their sequences, which differ by only 16 %.

The goal of our study was to identify the regions that mediate ACR-16 RIC-3 requirement using a series of chimera and point mutations, a commonly used technique when identifying functionally significant regions in receptors ^72–75^. Identifying the regions may help reveal how nascent subunits assemble in the absence of RIC-3 and/or the mechanistic details of what RIC-3 does when promoting subunit assembly ^69^. The high sequence similarity between the two genes allowed us to modify the *A. suum* and *D. medinensis acr-16* coding sequences to introduce restriction enzyme recognized sites without changing the protein product. No significant difference was measured between native and restriction modified sequences, nor their requirements for RIC-3, this was expected since RIC-3 effects on nematode receptors is at the protein level. Therefore, any changes in RIC-3 requirement would be due to the exchanged region. We identified multiple regions that affect the requirement for RIC-3 in oocytes using chimeric exchanges between the two subunits. The ICL and TM4 regions contributed marginally, with the contribution of the ICL apparently not localized, but distributed over the ICL. Two residues; I504 in the C-terminal tail and K/R159 in the cys-loop, had a major effect, with a third residue in the cys-loop, D157, also contributing to a lesser extent.

RIC-3 was first identified in *C. elegans* and has since been extensively studied for its involvement for both vertebrate and invertebrate receptors ^33, 37, 38^. It is predicted to have 1-2 transmembrane domains followed by an intracellular coiled-coil region and a disordered region^33, 47^. RIC-3 functions in the ER by binding directly to nascent subunits in a one-to-one ratio ^45, 76^.

Once bound, the intracellular coiled-coil region of RIC-3 participates in homotypic interactions that binds to another RIC-3 coiled-coil region (that is also bound to a subunit), promoting subunit dimerization. One RIC-3 dissociates from this intermediate complex and the process continues until a pentamer is formed ^45^. Thus, direct physical interactions to both the subunits and to other RIC-3 proteins are critical for assembly to occur ^32, 34, 43, 48, 49, 71, 77^. Other reports suggest RIC-3 may also inhibit certain subunit types from exiting the ER thereby altering subunit composition in heteromeric receptors, and that this is caused by decreased RIC-3 dissociation from the subunit ^34, 43, 76, 77^. RIC-3 is critical for receptor expression and coordinates associating subunits in the right stoichiometry. Interactions between RIC-3 and the subunits are vital for this process yet not all receptors require RIC-3, and it is not known what process it facilitates for assembly when it is required, or why some receptors require it.

The most notable finding presented here was that two amino acids mediated a RIC-3 requirement for AChRs. If the subunit had both the I504 in the C-terminal tail and K/R159 in the cys-loop, then it no longer required any RIC-3 for functional synthesis. Mutating either I504T or K/R159E conferred a RIC-3 requirement. Mutating D157N also had an effect, but to a lesser extent. The terminal tail and cys-loop are on the outside of the extracellular region of the receptor and previous reports found that these regions directly interact ^9, 63^. In an alpha7-5HT-3 chimeric receptor, interactions between the C-terminal tail and cys-loop lock the cys-loop in a mature conformation, masking ER retention signals, thus promoting cell-surface expression ^62^. Many heteromeric N-AChR receptor subunits contain the TM1 ER retention signal (PL(Y/F)(Y/F)xxN) ^13^, whereas the ACR-16 subunits studied here have the homomeric RRR motif ^12, 78^ which might be involved.

There is growing evidence for the involvement of the C-terminal tail in receptor expression. Deletions of the tail residues resulted in non-functional cationic ^16^ and anionic ^17^ channels, and it has been proposed that the more hydrophobic the tail is, the more it will interact with the ECD ^63^. In our case the isoleucine allowed the receptor to no longer require RIC-3, whereas mutation to threonine required RIC-3. The cys-loop motif is most known for its interactions with the TM2-TM3 linker during receptor gating ^6, 79^, but it has been implicated in receptor synthesis ^80, 81^. It undergoes large structural changes during subunit folding that are required for assembly, and inhibiting them prevents the joining of subunits to the receptor complex, thus ablating assembly at intermediate steps ^11^. It is not known what the precise mechanism of action is of the two identified residues here. However, combining our results with what is currently known about these regions in receptor expression, we propose that it is possible that the hydrophobic isoleucine at position 504 of the C-terminal tail allows the tail to readily interact with the cys-loop. This interaction may orient the K/R159 cys-loop residue in a conformation that makes key interactions with neighbouring regions that either 1) mask ER retention signals or 2) promote the cys-loop conformation required for receptor assembly and maturation. Mutating either of the residues prevents this interaction cascade from occurring, halting assembly and/or ER exit (Figure 5). To overcome this, RIC-3 physically interacts with the nascent subunit in such a way to either orient the tail towards the ECD or directly promote the cys-loop structural change itself. This would predict that RIC-3 interacts with the outermost region of the subunit to promote critical structural changes during assembly and may explain why the isoleucines in TM4 also contributed to RIC-3 requirement. It would also explain why the addition of RIC-3 increased responses of *D. medinensis* ACR-16 even if it was not absolutely required; their interaction might boost receptor synthesis that is already occurring.

**Figure 5.**
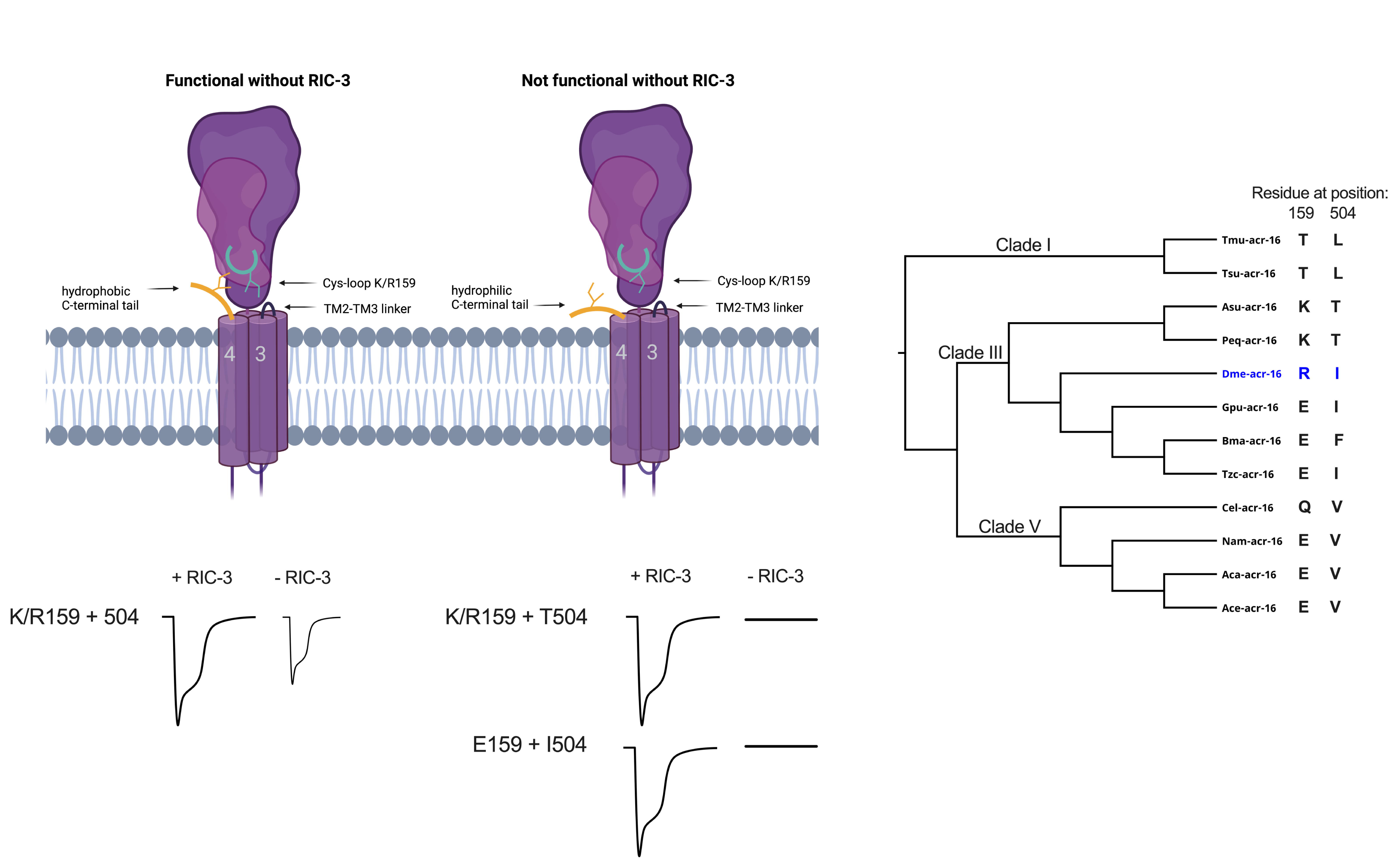
Proposed mechanism between C-terminal tail and cys-loop mediating RIC-3 requirement in acetylcholine receptors. When receptors have both cys-loop K/R159 and I504, function is possible in the absence of RIC-3. The hydrophobic I504 may orient the C-terminal tail towards the cys-loop in the ECD, this in turn orients the basic K/R159 cys-loop residue in a conformation that makes key interactions with neighbouring regions that either 1) mask ER retention signals or 2) promote the cys-loop conformation required for receptor assembly and maturation. Mutating either K/R159E or I504T prevents this interaction cascade from occurring, halting assembly and/or ER exit without RIC-3. To overcome this, RIC-3 physically interacts with the nascent subunit in such a way to either orient the tail towards the ECD or directly promote the cys-loop structural change itself.

No clearly stable interaction between residues at position 159 and 504 were seen in our molecular dynamic simulations, for either configurations that require RIC-3, or those that do not. This may be because the subunit model was based on a resolved structure of a mature, pentameric channel, i.e., a channel that has already gone through the structural changes that RIC-3 may affect. This structure shows the C-terminal tail as a helix projected away from the ECD ^66^. Now that these residues have been shown to influence the need for RIC-3, it will be important to assess whether they mediate a direct interaction with RIC-3, influence the ability of RIC-3 to perform its role or be directly involved in receptor synthesis. Intracellular localization of subunits that fail to be expressed at the surface may indicate which step in synthesis that is affected.

We also found that the ICL and TM4 contributed to RIC-3 requirement. This agrees with some, but not all, of the current literature. For the nematode receptors, the TM of RIC-3 was found to be required for the DEG/DES receptor ^43^, whereas only the intracellular coiled-coils were required for ACR-16 ^43, 44, 71^. However, for the mouse alpha7 receptor, the RIC-3 intracellular coiled-coil is not required ^69^. In a vertebrate chimeric receptor, a single residue near its TM1 was found to regulate RIC-3s inhibitory effects ^34^ while others found that the first 24 amino acids of the 5-HT_3_R ICL binds to RIC-3 ^50^. In the vertebrate AChRs, one study reported that the ECD and TMs interacts with RIC-3 ^69^. These discrepancies can be explained by the fact that RIC-3 is highly versatile and its function depends on the receptor type ^46^, the expression cell system ^40, 82^, its concentration ^70, 83, 84^ and its isoform ^31, 76, 85^. Additionally, the relative contribution of the different regions in RIC-3 are also subunit-dependent ^43, 44, 71^. Therefore, when subunits belong to different classes, they could be interacting with RIC-3 under different mechanisms that would explain RIC-3s versatility ^43, 44^. Some studies identify the regions for function while others identify the regions of interaction, and these may be not the same.

In addition to identifying the regions that mediated RIC-3 requirement, conclusions can be made on the observations that some chimeras had significantly different responses with RIC-3. First, chimeras that contained the *A. suum* ECD had higher responses compared to the same chimera that had the *D. medinensis* ECD (ex. Dme-ACR-16 vs Dme-asuECD, Asu-ACR-16 vs Asu-dmeECD, Dme-asuICL vs Asu-dmeTM; Dme-asuTM vs Asu-dmeICL). The *A. suum* ECD sequence appears to enhance the level of expression of the receptor, most likely at later stages since its effects are only observed if the receptor already being expressed. This coincides with the fact that the ECD contains many of the expression signal sequences ^15, 86–88^. Second, chimeras that split the ICL had decreased responses with *A. suum* TM4 (Asu-dmeICLa and Asu-dmeICLb). It is possible that sequences across the *A. suum* ICL and TM4 coevolved for efficient synthesis. However, the largely unknown structure and exact function of the ICL makes interpreting these results challenging. Some of the point mutations as well differed from their reference receptor (Figure S3A, Dme-TM4-3, Asu-TM4-1, Asu-TM4-1+2, Asu-TM4-2+3, Asu-tail, Asu-TM4-1+2+3+tail). These residues, along with the ECD and ICL, may provide an opportunity for further site-specific identification of regions critical for receptor expression.

Accessory proteins for pLGIC expression and regulation are becoming increasingly important. In order to understand the mechanisms by which this is determined, it is necessary to investigate residues of both accessory proteins and pLGIC subunits that affect this interaction. The observation that one species, out of twelve nematodes encoded an ACR-16 that could be expressed in *X. laevis* oocytes without RIC-3 suggested a discrete change within that species. This provided an opportunity to demonstrate that a single amino acid in the tail region was responsible for a majority of the effect. The fact that a substitution within the cys-loop was able to prevent, or enhance the requirement suggested that a physical interaction between the tail and cys-loop may be responsible. We have identified two residues that are critical in regulating AChR functional synthesis. Since they also determined receptor requirement for RIC-3, their interactions and function during synthesis may provide insight into the molecular mechanism of action that RIC-3 promotes. RIC-3 is encoded within the *D. medinensis* genome therefore it is unclear why this change has occurred in ACR-16 or what biological effect it has in the worm.

Uncovering the precise mechanism of how these two residues contribute to receptor expression may provide insight into how subunits and evolve requirements for accessory proteins and their biological significance.

## Materials and Methods

### Ethics Statement

All work involving animals was carried out under the McGill University Animal Use Protocol 2022–7758 authorized by the Animal Care Committee of the Office of Research Ethics and Compliance, McGill University.

### Sequence alignments, Restriction Enzyme Modified sequences and phylogeny

*A. suum* and *D. medinensis acr-16* coding sequences were obtained from their respective genomes available at WormBase ParaSite (v. WBPS13) ^61, 89^. Sequences were aligned in Geneious (v. 9.0.5, Biomatters Ltd) using the MAFFT plugin (v1.5.0) ^90^. This alignment was then used to modify both *acr-16* nucleotide sequences to introduce corresponding restriction enzyme recognition sites that do not change the amino acid sequence (termed ACR-16-REM). Eleven restriction enzyme sites were introduced into each acr-16 (Figure S1A), and these allowed for quick and efficient exchange of sequences between the two to generate chimera and point-mutation sequences. For the TM4 and C-terminal tail alignment (Figure 4A), *acr-16* sequences were obtained from the genomes of the following nematode species; *Ancylostoma caninum, Ancylostoma ceylanicum, Ascaris suum, Brugia malayi, Caenorhabditis elegans, Dracunculus medinensis, Gonglyonema pulchrum, Necator americanus, Parascaris equorum, Trichuris muris, Trichuris suis,* and *Thelazia callipaeda* ^89^. Sequences were translated and aligned in Geneious (v. 9.0.5, Biomatters Ltd) using the MAFFT plugin (v1.5.0). For the tree in Figure 5, a cladogram of the nematode species was made based on established phylogeny ^91^.

### Chimera and point mutation cloning

*B. malayi ric-3* was cloned previously from adult female cDNA ^61^. Native *acr-16* sequences were also obtained previously ^61^ and REM *acr-16* sequences were synthesized by Gene Universal (USA) in the pTD2 expression plasmid ^92^. Chimeras were made by digesting ACR-16-REM plasmids with the appropriate restriction enzymes (ThermoFisher Scientific, USA) (S Table 1) at 37°C for two hours followed by agarose gel purification (Zymo Research, USA) and overnight ligation at 4°C (T4 DNA ligase, Promega, USA). Ligated chimeras were transformed into DH5α (ThermoScientific, USA) and sequence verified (McLab, USA).

TM4 and C-terminal tail point mutations were made by digesting Asu-ACR-16-REM and Dme-ACR-16-REM with Bsu15I and ApaI (ThermoScientific, USA) followed by gel purification (Zymo Research, USA). These were then ligated to oligo inserts containing the TM4 and C-terminal tail point mutations (S Table 2). Oligo inserts were made following the Sigma oligo anneal protocol. The primers were suspended in annealing buffer (10 mM Tris, pH 7.5 - 8.0, 50 mM NaCl, 1 mM EDTA) to a concentration of 100 μM. Equal volumes of complementary primers were combined and annealed by heating at 95°C for 3 min followed by a gradual decline to 25°C over 45 minutes. Oligos were then ligated at 4°C overnight to the appropriate digested ACR-16-REM sequence (T4 ligase, Promega, USA) and transformed into DH5α (ThermoScientific, USA) and sequence verified (McLab, USA).

The cys-loop mutation constructs were made by digesting Dme-ACR-16-REM and the Asu-tail chimera with FastDigest XhoI and HindIII (ThermoScientific, USA). Digests were gel purified (Zymo Research, USA) and ligated to inserts containing the cys-loop point mutations (S Table 2), synthesized from Gene Universal (USA) at 4°C overnight (T4 DNA ligase, Promega, USA). Cys-loop mutation ligation products were transformed into DH5α (ThermoScientific, USA) and sequence verified (McLab, USA). All sequences were cloned in the pTD2 oocyte expression vector which contains the gene of interest flanked by 3’- and 5’- *X. laevis* beta-globin UTRs and a 3’ poly-A tail ^92^.

### In vitro RNA transcription

A PCR of all constructs using forward primer pTD2F (5’-TTGGCACCAAAATCAACGGG–3’) and reverse primer SP6 was carried out with SuperFi DNA Polymerase (ThermoFischer Scientific, USA) and agarose gel purified (Zymo Research, USA). These primers amplify the gene insert, the 3’ and 5’ *X. laevis* beta-globin UTRs and a 3’ poly-A tail. In vitro transcription of the PCR products using the mMESSAGE mMACHINE T7 kit (Ambion, USA) was performed following manufacturers protocol. cRNA was precipitated with lithium chloride and dissolved in RNase-free water and the concentration was determined using a Nanodrop spectrophotometer. The desired subunit and *ric-3* combinations were mixed and diluted with nuclease-free water to final injection concentrations of 250 ng/μL each.

### *Xenopus laevis* oocyte extraction and injection

*Xenopus laevis* were used in this study in accordance with the McGill University Animal Use Protocol 2022-7758. Adult female frogs (Xenopus1, USA) were housed in the Xenoplus Housing System (Technoplast, Italy). Frogs were anaesthetized in 0.15% MS-222, pH 7.3 (Sigma-Aldrich, USA) and an incision was made on the side of the abdomen and oocytes extracted. Incisions were sutured and the frogs placed in an isolation tank where they were regularly monitored for 1 week until return to the colony tank. Extracted oocytes were prepared according to standard protocol ^93^. Lobes were placed in a Ca^2+^-free OR2 solution (82 mM NaCl, 2 mM KCl, 1 mM MgCl_2_, 5 mM HEPES buffer, pH 7.3) and separated into clusters of <10 oocytes using fine tweezers. Oocytes were then incubated for 90 minutes at room temperature in Ca^2^-free OR2 supplemented with 10 mg/mL collagenase type II (Sigma-Aldrich, USA) and washed in Ca^2^-free OR2. Defolliculated oocytes were transferred to ND96 solution (NaCl 96 mM, KCl 2 mM, CaCl_2_ 1.8 mM, MgCl2 1 mM and HEPES 5 mM, pH 7.3) supplemented with 2.5 mM sodium pyruvate (Sigma-Aldrich, USA)^94^. For injection, glass capillaries were pulled into injection needles (World Precision Instruments, USA) and the tips clipped with tweezers. Needles were first back filled with oil followed by the cRNA injection mixture. Oocytes were injected with 50 nL (12.5ng per injected gene) of cRNA mixture using a Nanoject injector (Drummond Scientific, USA). After injection, oocytes were incubated at 18°C in ND96 solution for 2 days until electrophysiology.

### Two-electrode voltage-clamp electrophysiology

Responses were measured after two days following cRNA injection. Oocytes were individually submerged in Ringer recording solution (NaCl 100 mM, KCl 2.5 mM, CaCl2 1 mM, HEPES 5 mM, pH 7.3) in an RC3Z chamber (Harvard Apparatus, USA) that was connected to a perfusion system containing Ringer recording solution and ligand solution. Pulled glass capillaries (World Precision Instruments, USA) were filled with 3M KCl and the tips clipped with tweezers and checked for appropriate resistance between 0.5 and 5 MΩ. A 3M KCl agar bridge was used to ground the oocyte bath chamber. Measurements were made using a Geneclamp 500B amplifier with Digidata1322M (Axon instruments, USA). Oocytes were voltage-clamped at -60 mV and any that had holding currents less than -400 nA or that had currents that did not stabilize were discarded due to poor oocyte condition. Current was measured in response to 100 µM ACh (Sigma-Aldrich, USA) dissolved in recording solution, pH 7.3. ACh ligand was applied with an approximate flow rate of 1.2 m/s and oocyte current responses measured. Oocytes were exposed to the ligand solution until a distinct current peak was observed followed by a wash with the recording solution until current returned to baseline. For examples of oocyte responses, recordings were filtered in Clampfit using the lowband pass at 40 Hz cutoff. Data was analyzed using Clampex 9.2 (Axon Instruments, USA) and graphs made in Prism (version 9.0) with significance determined using t-tests and error bars representing standard error.

### Molecular dynamics

Homology models of Asu-ACR-16 and Dme-ACR-16 were prepared using the default hm_build.mcr macro of YASARA (v.16.3.8, YASARA Biosciences) ^67^ based on the human a modified alpha4 subunit template (PDB:6CNK-A) ^95^ after removing the signal peptide and intracellular loop. A terminal sequence YIIA was added to the template and the tail energy minimized so that the residue equivalent to I504 was 5 Å from the cys-loop residues equivalent to R/K159. Each model was the starting point for a membrane embedded molecular dynamics simulation for 10 ns, using the default md_runmembranefast.mcr macro. Simulations were enabled by support provided by CalculQuebec (https://www.calculquebec.ca/en/) and Compute Canada (www.computecanada.ca).

## Description of supplementary material

Supplemental Figure 1 (FigureS1) *A. suum* and *D. medinensis* acr-16 sequence alignment. This figure shows the alignment between the genome predicted *D. medinensis* and *A. suum* ACR-16 sequence and the Restriction Enzyme Modified ACR-16 coding sequences used in the present study as well as the protein alignment.

Supplemental Figure 2 (FigureS2). Responses of *A. suum* and *D. medinensis* ACR-16 chimeras with and without RIC-3. This figure contains the raw currents recorded from the chimeras, both with and without RIC-3.

Supplemental Figure 3 (FigureS3). Responses of *A. suum* and *D. medinensis* ACR-16 TM4 and C-terminal tail point mutation mutants with and without RIC-3. This figure contains the raw currents recorded from the TM4 point mutations, both with and without RIC-3.

Supplemental Table 1. Restriction enzymes used to generate the chimeras. This table contains the restriction enzymes and plasmids used to generate the chimeras.

Supplemental Table 2. *A. suum* and *D. medinensis* TM4 point mutation primers and cys-loop insert sequences. This table contains the primers used to generate the TM4 point mutations and the inserts containing the cys-loop mutations.

## Author Statement

Nicotinic channels often require the RIC-3 accessory protein for function in vivo and in heterologous expression systems. However, not all channels require RIC-3 and even closely related receptors can have differing requirements for it. Using two closely related receptors with opposite RIC-3 requirements, here we identify two residues in the extracellular region of the receptor that determine RIC-3 requirement. We propose that these residues participate in critical interactions that either promote receptor expression without RIC-3 or directly interact with it.

## Author contributions

Conceptualization, Visualization, Methodology & Manuscript editing – JDN & RNB

Investigation, Manuscript writing & Formal analysis - JDN

Funding Acquisition & Resources – RNB

## Abbreviations

5-HT3R: Serotonin receptor
ACh: Acetylcholine
AChR: Acetylcholine receptor
ACR-16: Acetylcholine receptor 16
ACR-16-REM: Restriction enzyme modified ACR-16
Asu: *Ascaris suum*
Asu-ACR-16: *Ascaris suum* ACR-16 receptor
cRNA: complimentary RNA
Dme: *Dracunculus medinensis*
Dme-ACR-16: *Dracunculus medinensis* ACR-16 receptor
ECD: Extracellular domain
ER: endoplasmic reticulum
GABA: γ-aminobutyric acid
ICL: Intracellular loop
N-AChR: Nicotinic acetylcholine receptor
pLGIC: pentameric ligand-gated ion channel
RIC-3: Resistance to cholinesterase 3
TM: Transmembrane domain

## Acknowledgements

The funding for this research was a Discovery Research Grant (RGPIN/05320-2020) from the Canadian National Sciences and Engineering Research Council to RNB. JDN was the recipient of an NSERC PGSD Scholarship.

## Supplemental Information

**Figure S1. *A. suum* and *D. medinensis* acr-16 sequence alignment**.

**(A)** Nucleotide alignment of genome-predicted Asu-ACR-16 and Dme-acr-16 along with the restriction enzyme modified (REM) sequences.

**(B)** Protein sequence alignments for *A. suum* and *D. medinensis* ACR-16 show widespread conservation with majority of substitutions observed in the intracellular loop. Three substitutions are observed in the TM4, one in the C-terminal tail, and one in the cys-loop. Cys-loop shown in red, transmembrane domains shown in green.

**Figure S2. Responses of *A. suum* and *D. medinensis* ACR-16 chimeras with and without RIC-3.**

Chimeras were made exchanging regions between *A. suum* and *D. medinensis* ACR-16 to identify the residues determining RIC-3 requirement. Responses to 100 µM ACh are shown.

**(A)** Chimera response with RIC-3 co-injection. Comparing responses with RIC-3 to Asu-ACR-16-REM (2197 ± 85 nA), Asu-dmeICL (1418 ± 87 nA, p<0.0001) and Asu-dmeECD (1779 ± 69 nA, p<0.01), had significantly smaller responses whereas Asu-dmeTM (2415 ± 111 nA, p=0.1269), was not statistically different. For the TM2/4 chimeras, Asu-dmeICL+TM2 (1528 ± 59 nA, p<0.0001) had lower responses and Asu-dmeICL-TM4 (2592 ± 60 nA, p<0.01) had higher responses. Both chimeras that split the ICL in half produced lower responses; Asu-dmeICLa (1104 ± 72 nA, p<0.0001) and Asu+dmeICLb (908 ± 130 nA, p<0.0001). For the *D. medinensis* chimeras, compared to Dme-ACR-16-REM (1989 ± 73 nA), Dme-asuECD (2242 ± 62 nA, p<0.05) had significantly larger responses whereas Dme-asuICL (2233 ± 106 nA, p=0.0576) and Dme-asuTM (1979 ± 104 nA, p=0.9421) were not different. Therefore the ECD may enhance the functional expression of receptors even in the presence of RIC-3. For the TM2/4 chimeras, Dme-asuICL+TM4 (2123 ± 107 nA, p=0.2939) had comparable responses whereas Dme-asuICL+TM2 (3110 ± 85nA, p<0.0001) had higher responses. The chimeras that split the ICL in half produce responses that did not differ from Dme-ACR-16, Dme-asuICLa (2183 ± 66 nA, p=0.0601) and Dme-asuICLb (2036 ± 82 nA, p=0.6727).

**(B)** Chimera responses without RIC-3 co-injection. For the *A. suum* chimeras, comparing responses to Asu-ACR-16-REM (13.92 ± 4.3 nA), Asu-dmeICL (211.2 ± 24.16 nA, p<0.0001) and Asu-dmeTM (1064 ± 93.6, nA p<0.0001) had significantly larger responses whereas Asu-dmeECD (0.0 ± 0 nA, p=0.01), was lower. For the TM2/4 chimeras, Asu-dmeICL+TM2 (89.5 ± 11 nA, p<0.0001) and Asu-dmeICL+TM4 (2454 ± 72 nA, p<0.0001) had larger responses. Both chimeras that split the ICL in half produced no response; Asu-dmeICLa (0 ± 0 nA, p<0.0001) and Asu+dmeICLb (0 ± 0 nA, p<0.0001). For the *D. medinensis* chimeras, compared to Dme-ACR-16-REM (1047 ± 60.6 nA), Dme-asuECD (1563 ± 102nA, p<0.0005) had significantly larger responses whereas Dme-asuICL (251.3 ± 30.5 nA, p < 0.0001) and Dme-asuTM (43.55 ± 8.81 nA, p<0.0001) were lower. For the TM2/4 chimeras, Dme-asuICL+TM2 (1527 ± 103 nA, p<0.0005) was larger whereas Dme-asuICL+TM4 (0 ± 0 nA, p<0.0001) had no response. The chimeras that split the ICL in half produced lower responses, Dme-asuICLa (575 ± 81 nA, p<0.0001) and Dme-asuICLb (383 ± 45 nA, p<0.0001). Error represents standard error.

*p<0.05; **p<0.01; ***p<0.0005; ****p<0.0001.

**Figure S3. Responses of *A. suum* and *D. medinensis* ACR-16 TM4 and C-terminal tail point mutation mutants with and without RIC-3**.

Point mutations were made in *A. suum* and *D. medinensis* ACR-16 in the fourth transmembrane domain and/or C-terminal tail. Current responses to 100 µM ACh are shown. All TM4 and C-terminal tail point mutation produced robust responses with RIC-3 with some significant deviations from their reference receptors’ responses.

**(A)** For the mutations within Asu-ACR-16 when including RIC-3, three conditions produced statistically smaller responses Asu-TM4-1 (1952 ± 74 nA, p<0.05), Asu-TM4-1+2 (1851 ± 139 nA, p<0.05) and Asu-TM4-2+3 (1666 ± 76 nA, p<0.0001), and both tail mutation conditions produced statistically larger responses; Asu-TM4-1+2+3+tail (3070 ± 106 nA, p<0.0001) and Asu-tail (2560 ± 60 nA, p<0.01). Whereas mutations to Asu-TM4-2 (1923 ± 109 nA, p=0.0542), Asu-TM4-3 (2309 ± 60 nA, p=0.3373), Asu-TM4-1+3 (2338 ± 73 nA, p=0.2566) and Asu-TM4-1+2+3 (1977 ± 95 nA, p=0.1035) were not different.

**(B)** Only one of the point mutations within Dme-ACR-16 produced significantly different responses when including RIC-3. The single point mutation to the Dme-TM4-3 had lower responses (1710 ± 72 nA, p<0.05) while all others produced responses indistinguishable from Dme-ACR-16-REM; Dme-TM4-1 (2100 ± 77 nA, p=0.3102), Dme-TM4-2 (1776 ± 101nA, p=0.0889), Dme-TM4-1+2 (1987 ± 106 nA, p=0.9911), Dme-TM4-1+3 (2033 ± 86 nA, p=0.6971), Dme-TM4-2+3 (2074 ± 124 nA, p=0.5382), Dme-TM4-1+2+3 (1925 ± 47 nA, p=0.4840), Dme-TM4-1+2+3+tail (2169 ± 111 nA, p=0.5539), Dme-tail (1920 ± 90 nA, p=0.1657).

**(C)** For the mutations within Asu-ACR-16 without RIC-3, three conditions were not statistically different from Asu-ACR-16; ASU-TM4-1 (31 ± 7 nA, p=0.0556), ASU-TM4-2 (23 ± 4 nA, p=0.1250) and Asu-TM4-2+3 (24 ± 3 nA, p=0.0627). All other mutation conditions produced significantly higher responses without RIC-3 that were still minimal; Asu-TM4-3 (81 ± 18 nA, p<0.01), Asu-TM4-1+2 (49 ± 7 nA, p<0.0005), Asu-TM4-1+3 (102 ± 16 nA, p<0.0001), and Asu-TM4-1+2+3 (134 ± 29 nA, p<0.0005). Asu-tail (1659 ± 84 nA, p<0.0001) and Asu-TM4-1+2+3+tail (2914 ± 70 nA, p<0.0001) produced the largest responses.

**(D)** For the mutations within Dme-ACR-16 without RIC-3, all conditions were statistically smaller from Dme-ACR-16-REM. Dme-TM4-1 (573 ± 59 nA, p<0.0001), Dme-TM4-2 (843 ± 40 nA, p<0.05), Dme-TM4-3 (591 ± 65 nA, p<0.0001), Dme-TM4-1+2 (616 ± 43 nA, p<0.0001), Dme-TM4-1+3 (420 ± 43 nA, p<0.0001), Dme-TM4-2+3 (607 ± 49 nA, p<0.0001) and Dme-TM4-1+2+3 (799 ± 54 nA, p<0.01) produced slightly smaller responses whereas Dme-tail (51 ± 7 nA, p<0.0001) and Dme-TM4-1+2+3+tail (33 ± 10 nA, p<0.0001) has the most inhibited responses.

**Table S1.** Restriction enzymes used to generate the chimeras. ACR-16 chimeras (Figure 1) were made by digesting the ACR-16-REM sequences (S Figure 1) using the designated restriction enzymes.

|  | “Background” digestion | “Insert” Digestion | Restriction enzymes |
| --- | --- | --- | --- |
| Dme-asuICL | DME-ACR-16-REM | ASU-ACR-16-REM | PspLI + ClaI |
| Asu-dmeICL | ASU-ACR-16-REM | DME-ACR-16-REM | PspLI + ClaI |
| Dme-asuECD | DME-ACR-16-REM | ASU-ACR-16-REM | NotI + AccIII |
| Asu-dmeECD | ASU-ACR-16-REM | DME-ACR-16-REM | NotI + AccIII |
| Dme-asuTM | ASU-dmeECD | DME-asuECD | PspLI + ClaI |
| Asu-dmeTM | DME-asuECD | ASU-dmeECD | PspLI + ClaI |
| Asu-dmeICLa | DME-asuICL | ASU-dmeICL | NotI + SfuI |
| Asu-dmeICLb | ASU-ACR-16-REM | DME-ACR-16-REM | NotI + SfuI |
| Dme-asuICLa | DME-ACR-16-REM | DME-asuICL | NotI + SfuI |
| Dme-asuICLb | DME-asuICL | DME-ACR-16-REM | NotI + SfuI |
| Asu-dmeICL+TM2 | ASU-ACR-16 | DME-ACR-16-REM | AccIII + ClaI |
| Asu-dmeICL+TM4 | ASU-dmeTM | DME-asuTM | AccIII + ClaI |
| Dme-asuICL+TM2 | DME-asuTM | ASU-dmeTM | AccIII + ClaI |
| Dme-asuICL+TM4 | DME-ACR-16 | ASU-ACR-16 | AccIII + ClaI |

**Table S2.**
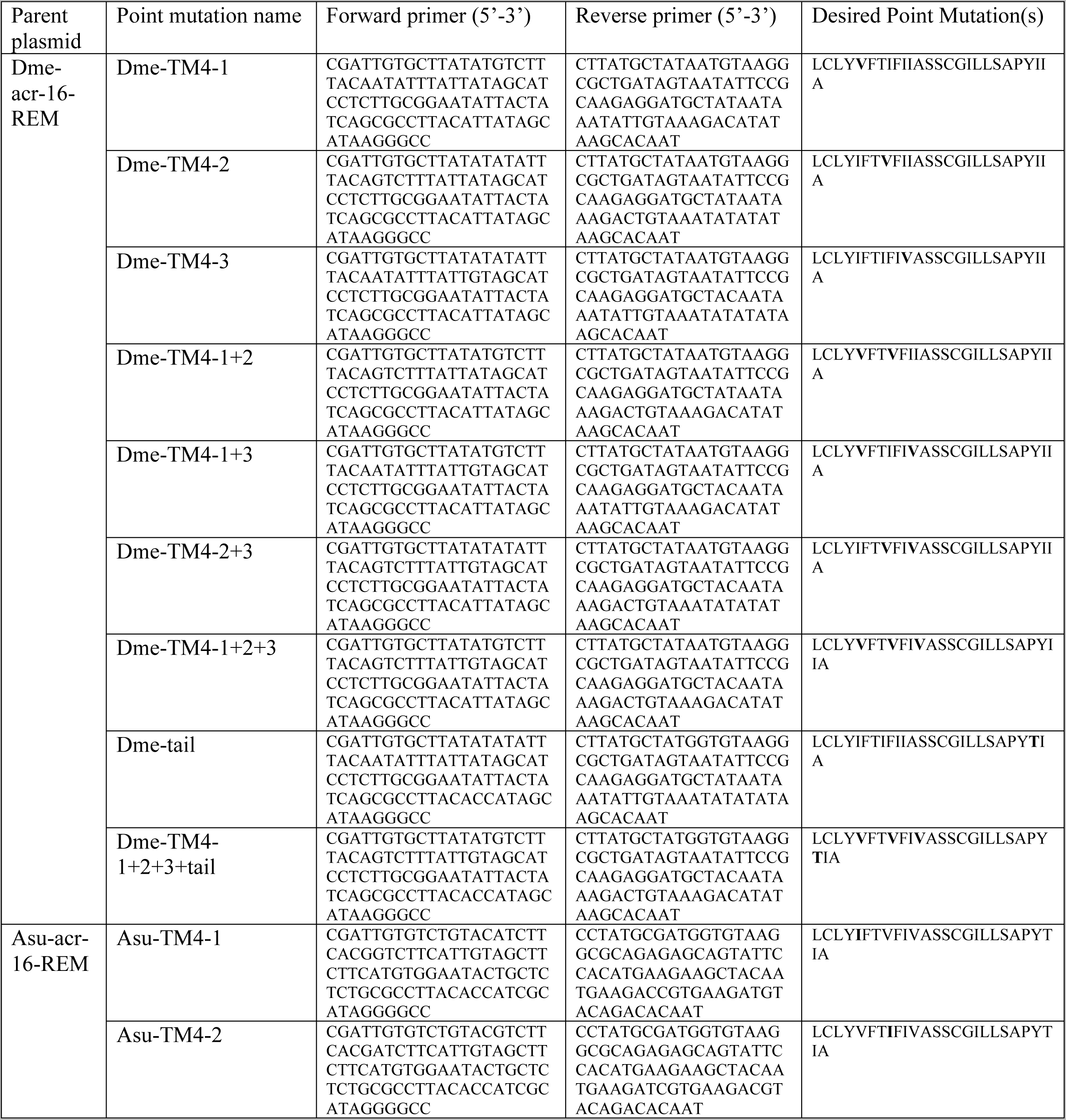
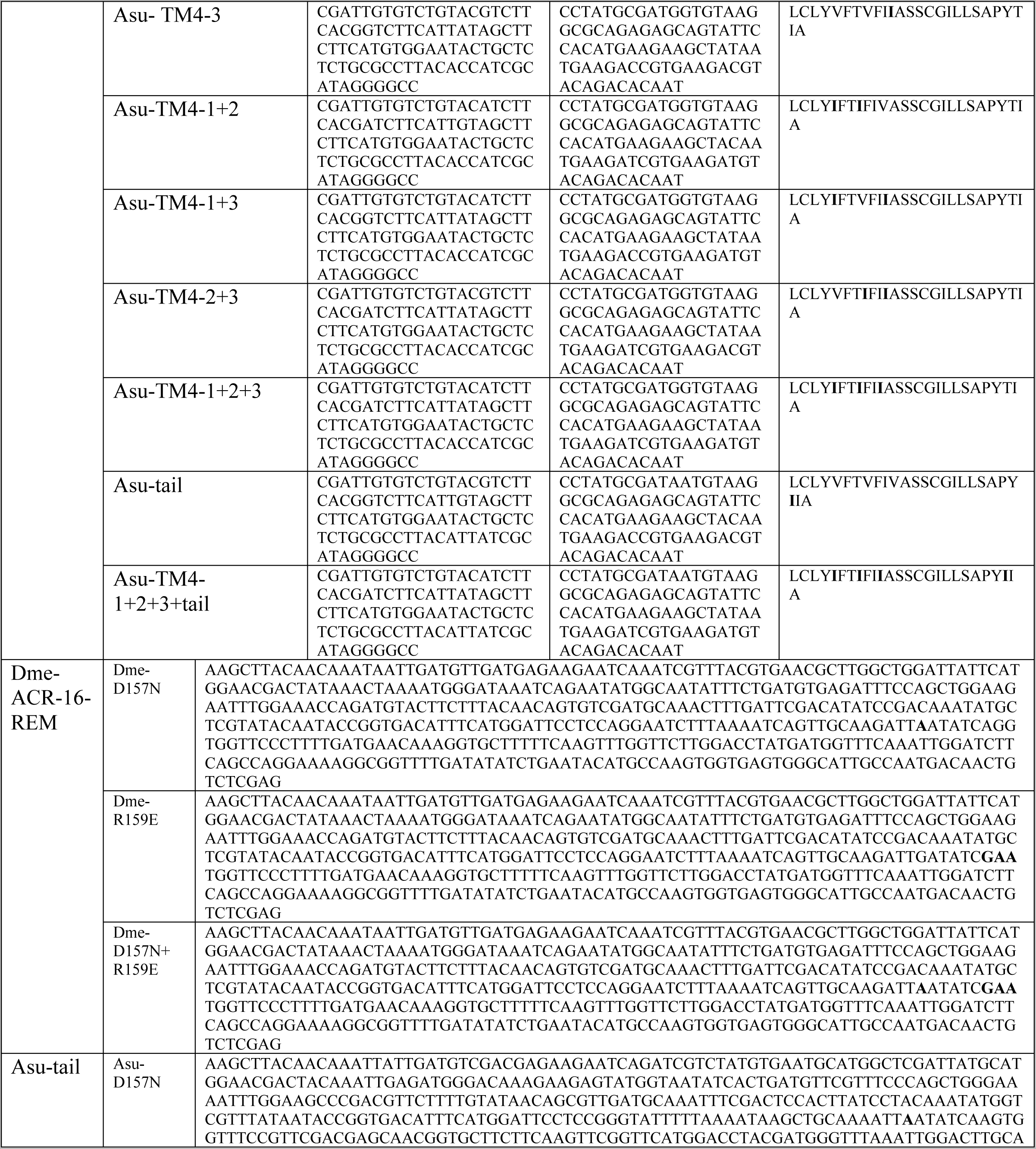
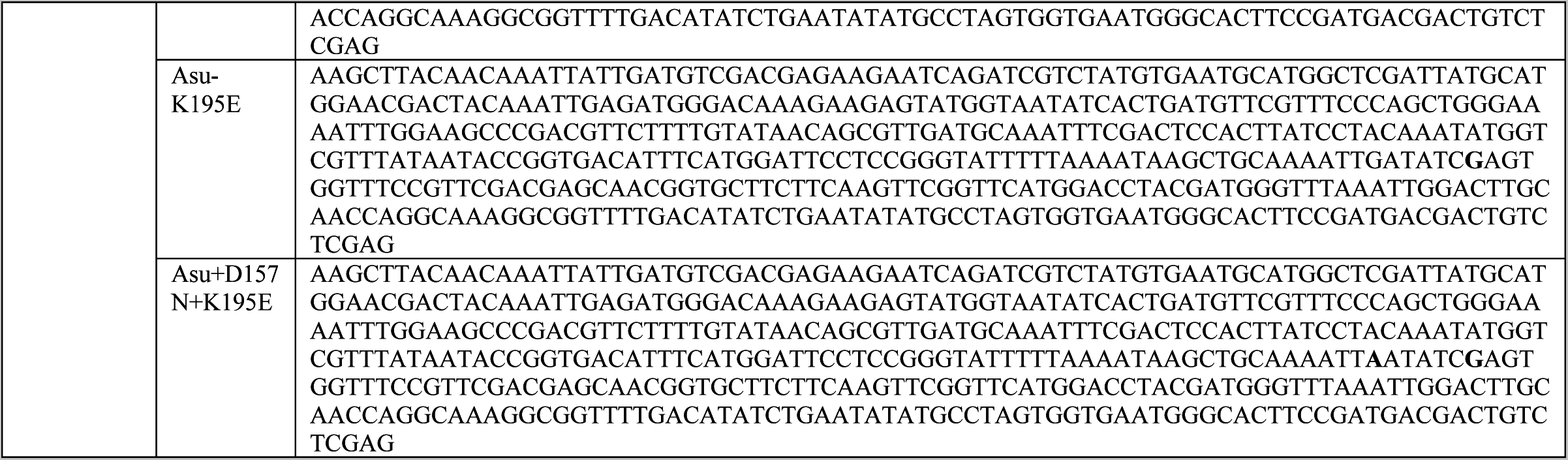
*A. suum* and *D. medinensis* TM4 point mutation primers and cys-loop insert sequences. For the TM4 and C-terminal tail point mutations, point mutations were introduced into the fourth transmembrane domain and C-terminal tail of *A. suum* and *D. medinensis* ACR-16. Desired point mutations are shown with the mutated residues in bold. For the cys-loop point mutations, Dme-ACR-16-REM and Asu-tail were digested with HindIII and XhoI followed by ligation to inserts containing the point mutations. The nucleotide sequence of the cys-loop point mutation inserts are shown with the mutations in bold.

