## Supplementary figures and images for "Two residues determine nicotinic acetylcholine receptor requirement for RIC-3"

### Supplemental Figure 1

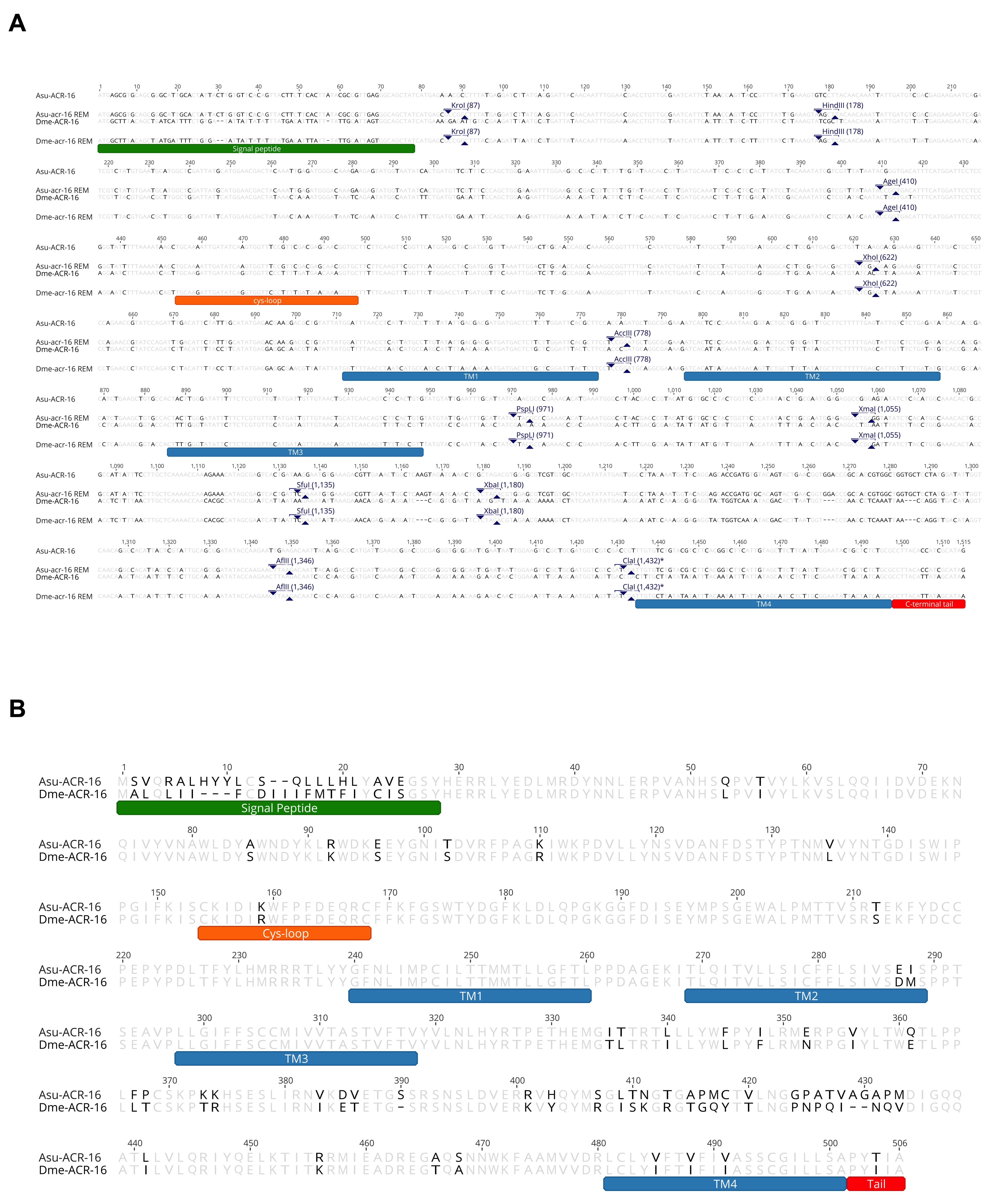

### Supplemental Figure 2

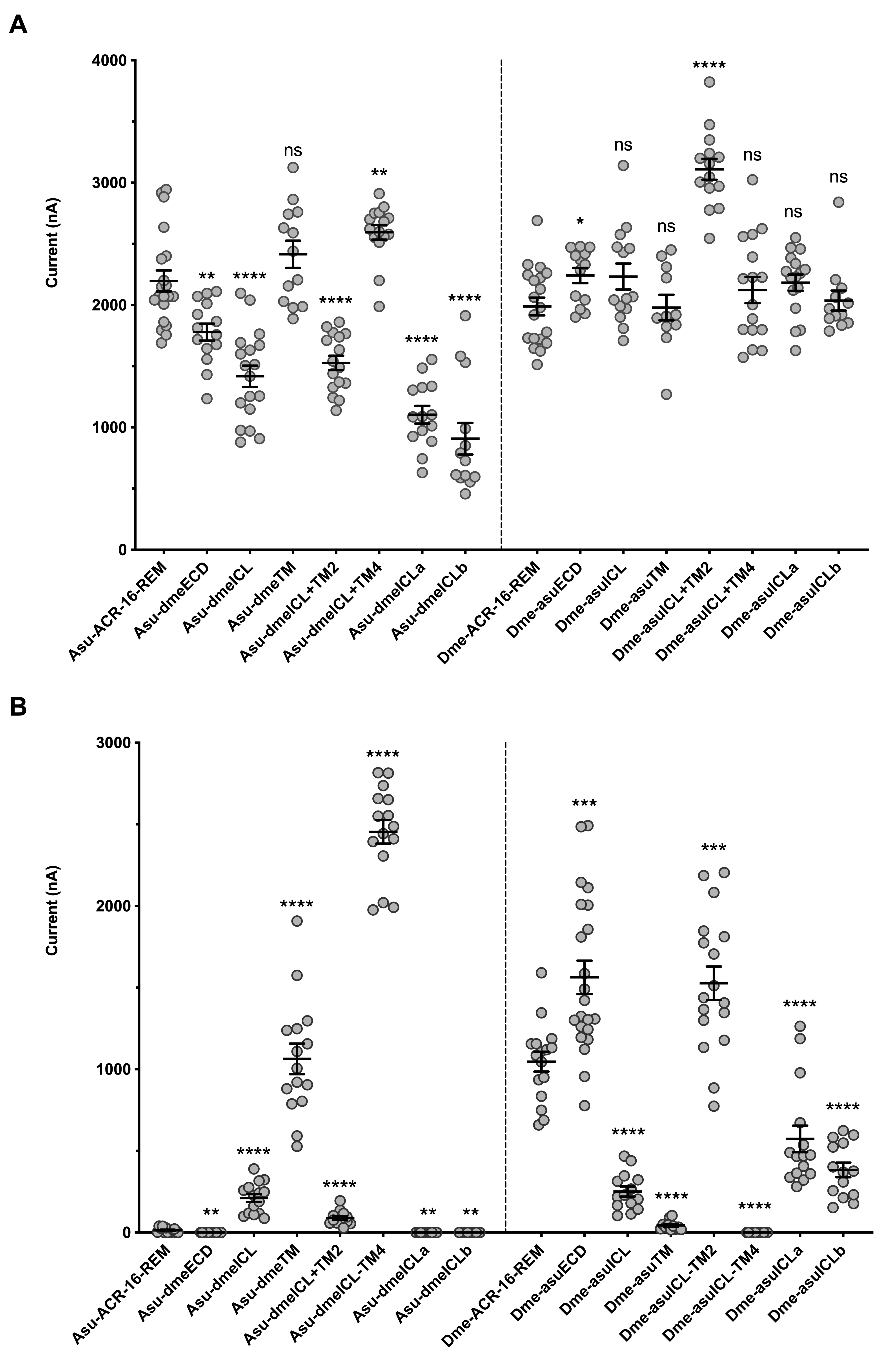

### Supplemental Figure 3

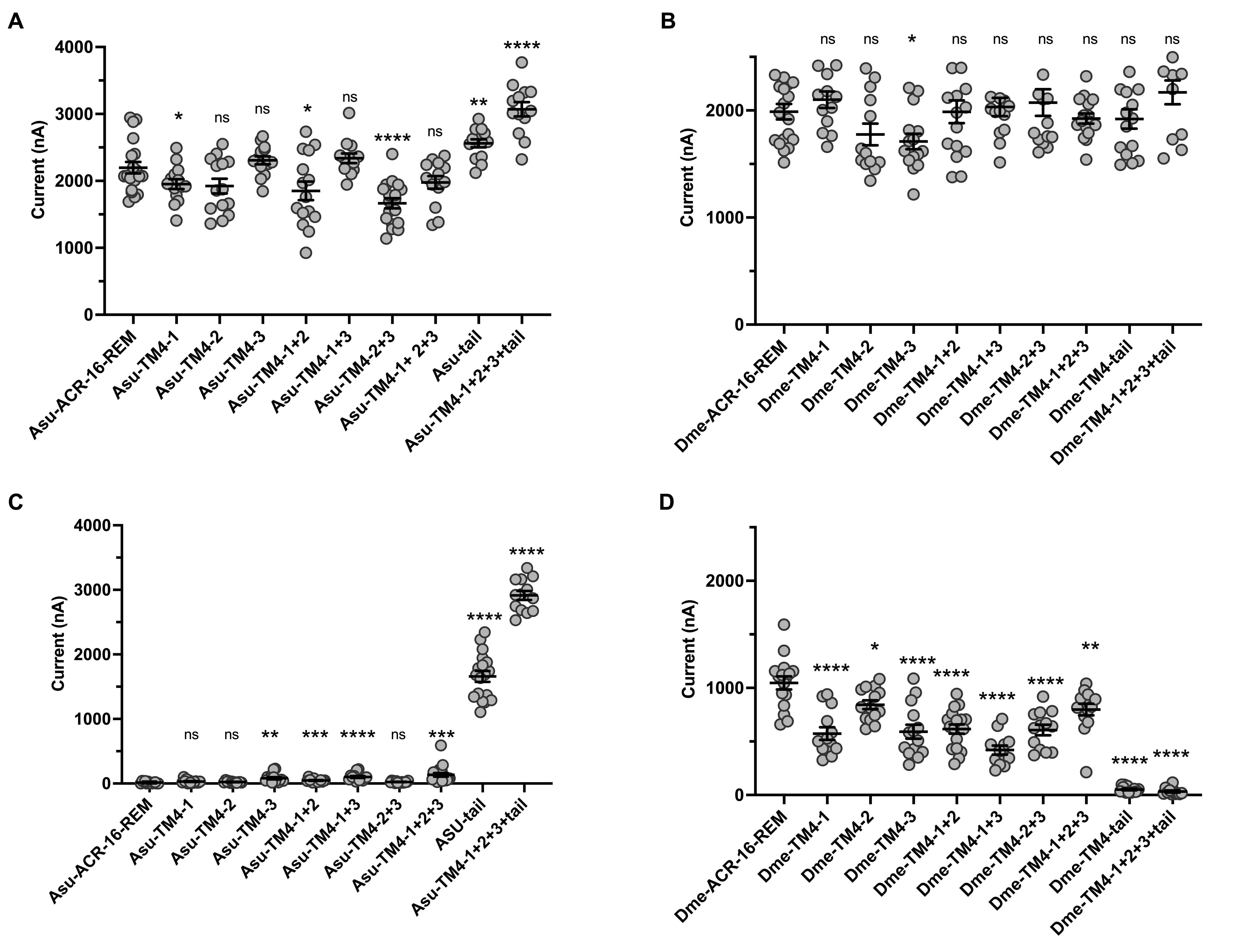
